# SCO-spondin knockout mice exhibit small brain ventricles and mild spine deformation

**DOI:** 10.1101/2023.08.01.551512

**Authors:** Huixin Xu, Guillaume P. Dugué, Yasmine Cantaut-Belarif, François-Xavier Lejeune, Suhasini Gupta, Claire Wyart, Maria K. Lehtinen

## Abstract

Reissner’s fiber (RF) is an extracellular polymer comprising the large monomeric protein SCO-spondin (SSPO) secreted by the subcommissural organ (SCO) that extends through cerebrospinal fluid (CSF)-filled ventricles into the central canal of the spinal cord. In zebrafish, RF and CSF-contacting neurons (CSF-cNs) form an axial sensory system that detects spinal curvature, instructs morphogenesis of the body axis, and enables proper alignment of the spine. In mammalian models, RF has been implicated in CSF circulation. However, challenges in manipulating *Sspo*, an exceptionally large gene of 15,719 nucleotides, with traditional approaches has limited progress. Here, we generated a *Sspo* knockout mouse model using CRISPR/Cas9-mediated genome-editing. *Sspo* knockout mice lacked RF-positive material in the SCO and fibrillar condensates in the brain ventricles. Remarkably, *Sspo* knockout brain ventricle sizes were reduced compared to littermate controls. Minor defects in thoracic spine curvature were detected in *Sspo* knockouts, which did not alter basic motor behaviors tested. Altogether, our work in mouse demonstrates that SSPO and RF regulate ventricle size during development but only moderately impact spine geometry.

## Introduction

The subcommissural organ (SCO) is a circumventricular organ that shares some functions with the choroid plexus (ChP), the principal source of cerebrospinal fluid (CSF) and a blood-CSF barrier. Like the ChP, the SCO secretes a range of regulatory proteins into the CSF including growth factors and binding proteins (e.g., Transthyretin) that may contribute to CNS development [1]. The SCO is also hypothesized to regulate ventricular volume like the ChP [2], but our understanding of SCO functions is very limited.

In adult organisms the SCO is the sole source of a unique member of the spondin family of proteins termed SCO-spondin (SSPO), a gigantic glycoprotein (close to 5,000 amino acids, see [3]). Soluble monomeric SSPO can fold in different configurations and its numerous motifs enable interactions with diverse signaling molecules in the CSF including monoamines, growth factors, and lipids [4, 5]. Like other so-called matricellular proteins, SSPO is likely to have a diversity of functions mirroring its structural complexity. In addition to maintenance of CSF flow dynamics, proposed functions of SSPO include neurogenesis and neuroprotection [6–9]. SSPO may also serve to regulate bioavailability of substances in the CSF when its binding capacity is sufficient to limit the concentration of unbound ligands. In addition to its soluble monomeric form, SSPO can polymerize to form Reissner’s fiber (RF), a long extracellular threadlike protein aggregate that emerges during embryogenesis and runs from the SCO through the cerebral aqueduct, the fourth ventricle, and into central canal of the spinal cord [10]. Despite its presence across numerous vertebrate species, the function of the RF in mammals is unclear. However, there is good evidence that the RF plays a critical role in developing zebrafish [11]. Together with CSF-contacting neurons (CSF-cNs), the RF forms an axial sensory system that has been shown in zebrafish to detect spinal curvature, instructs morphogenesis of the body axis, and enables proper alignment of the spine [12, 13] [58].

Here, we leverage a *Sspo* knockout (*Sspo*^-/-^) mouse that we generated using CRISPR/Cas9-mediated genome editing to explore the functions of SSPO in mice by determining the effects of loss of SSPO on ventricular volume, spine curvature and motor function. We found that the brain ventricle volumes were reduced in *Sspo*^-/-^ mice compared to their wild type (*Sspo*^+/+^) siblings, without defects in the central canal. Unlike studies in zebrafish, we observed only minor defects in spine geometry at the thoracic level and no differences in performance for basic motor assays. Altogether, our work demonstrates in mice that SSPO and RF are involved in regulating the volume of the CSF during development but loss of RF has only moderate impact on spine geometry.

## Methods

### Animals and genome editing

All experiments involving mice in this study were approved by the BCH IACUC. Germline *Sspo* knockout (*Sspo^tm1^* -/-) mice were generated using genome editing on a C57BL/6 background. Animals were housed in a temperature-controlled room on a 12-hr light/12-hr dark cycle and had free access to food and water. Comparable number of males and females were included. *Sspo*^+/+^ and *Sspo*^-/-^ mice were littermates generated by breeding heterozygous males and females.

*Sspo*^-/-^ mice were generated using the following guide RNA:

Sspo-Ex2-1 TCCATAGCATCCCAAAGAGA
Sspo-Ex3-1 GGTACGAGGTCCTCCTGACG
Sspo-Ex4-1 GTAGCAGGCCTGGTTTCCGG
Sspo-Ex6-1 CAGCTAGCAAGTAGGTACAC

Progenies were screened by loss of restriction enzyme sites for successful modification and sequenced to determine the consequences of genetic modifications. Of all the mice screened, one founder carried a 5 bp deletion in exon 2, leading to an early stop codon. The founder was used to establish the *Sspo^tm1^* -/-colony. Subsequent genotyping was conducted using the following primer set: CTGTGAAGGGATGCTGGAGGTAG (forward) and GTACTATGAATCCGAGGCCCCAA (reverse). The product from wild type mice is a 426bp band that can be cut by HpyAV. Mice carrying the 5bp deletion will yield a band that cannot be digested. Later genotyping was conducted at Transnetyx.

### Immunohistochemistry

Mice were perfused intracardially with ice-cold PBS followed by cold 4% PFA. Brains were dissected, soaked in 4% PFA at 4°C overnight and cryopreserved in 30% sucrose for 2 days before snap-freezing in liquid nitrogen-cooled isopentane. Coronally oriented cryosections were prepared at 16 µm thickness using a CM3050S cryostat (Leica). Blocking was performed by incubating the sections in 3% BSA and 0.1% Tween-20 in PBS one hour at room temperature. Sections were then incubated overnight at 4°C with the L1P1b rabbit anti-serum (kind gift of S. Gobron; 1:250 dilution) [14] to detect RF-positive material in the SCO and ventricular space. After washing with 0.1% Tween-20 in PBS, sections were incubated with a Goat-anti-Rabbit-Alexa488 secondary antibody (Invitrogen; 1:400 dilution) for 45 minutes at room temperature with counterstaining by 4’,6-diamino-2-phenylindole (DAPI) to label nuclei. Sections were imaged on an inverted SP8 DLS confocal microscope (Leica) equipped with a 40X objective (oil immersion; NA = 1.3). The same imaging parameters (laser power, scanning speed, wavelength gating for photon collection) were used to image sections derived from wild type and knockout animals. Images were processed using Fiji [15]. Maximal Z-projections of 10 µm in depth are represented (**Fig. 1**).

**Fig. 1.**
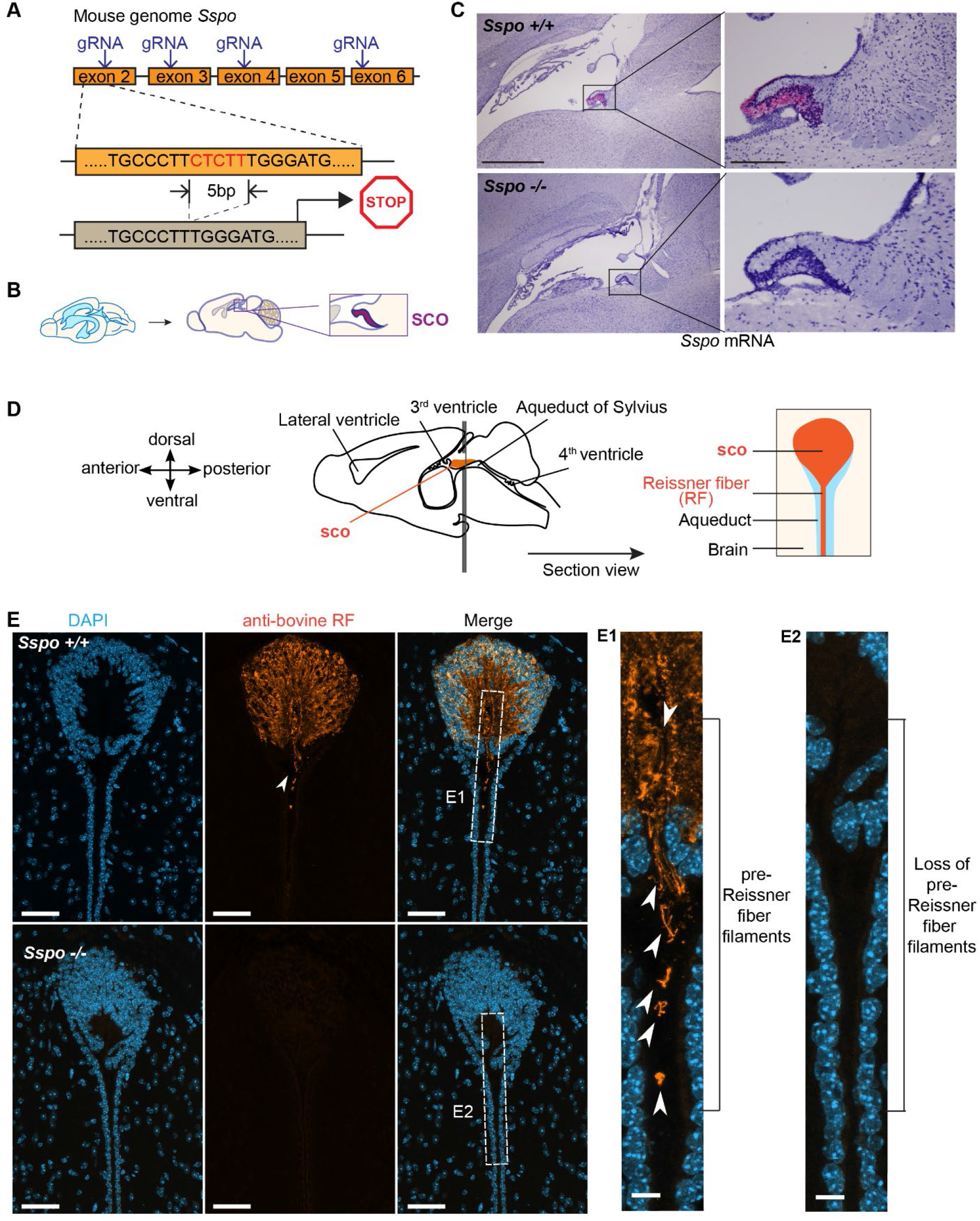
*Sspo*^-/-^ mice lack RF positive material in the SCO and ventricles. (**A**) Schematic depicting gene editing strategy. (**B**) Schematic depicting SCO location in sagittal brain sections corresponding to C. (**C**) BaseScope reveals lack of *Sspo* transcripts in the SCO of *Sspo*^-/-^ mice. Scale = 1mm (left panels) and 200 µm (right panels). (**D**) Schematic depicting the brain sectioning strategy taken to visualize SCO ependyma and proximal ventricular region. (**E**) RF positive material was not detected in *Sspo*^-/-^ mice vs. controls. Scale = 50 µm.

### BaseScope

Mouse brain sections were prepared as described above in RNAse-free conditions and stored at -80°C. BaseScope (ACD Bio, 323910) was used to detect the presence or absence of *Sspo* mRNA. A customized probe (1zz probe named BA-Mm-sspo-1zz-st, targeting 45-85 of NM_173428.4 *Sspo* sequence) was designed to distinguish wild type sequence vs. mutant sequence with 5bp deletion.

### Magnetic resonance imaging (MRI)

Mice were imaged using Bruker BioSpec small animal MRI (7T) at 6 weeks while under anesthesia by isoflurane. A warm pad was used to maintain body temperature. Breathing rate and heart rate were monitored to reflect the depth of anesthesia. All axial T2 images were acquired using the following criteria: TE/TR=60/4000; Ave=8; RARE=4; slice thickness = 0.6mm.

### Calculation of the ventricular volume

Ventricle volumes were calculated by manual segmentation using FIJI/ImageJ. Lateral ventricle and 3rd ventricle areas, which appear white in T2 MR images, were hand selected to measure total area by FIJI. Ventricle CSF volume was calculated by total ventricle area × 0.6 mm slice thickness. Data were blinded during analysis to avoid bias – see also [16].

### Behavioral analysis

All behavioral tests were conducted by BCH Animal Behavior and Physiology Core. All operators of the tests were blinded to the IDs of the mice.

### Gripping test

Gripping test was performed using BIO-GS4 system to objectively quantify the muscular strength of mice. Briefly, the test determines the maximal peak force generated by a mouse when the operator tries to pull it out of a specially designed grid. Forelimb gripping strengths were tested in this study.

### Rotarod

Rotarod was performed using the IITC Rotarod system. Mice were placed on a rotating beam (1 ¼ inches diameter) with increasing rotation frequency, ranging from 5 r.p.m. to 40 r.p.m. over the course of 5 minutes. The length of time (seconds) each mouse stayed on the beam was recorded to reflect its ability of motor coordination. Each mouse was tested in 6 independent trials.

### Digigait

Mice were placed on a Digigait device which resembles a treadmill. A series of parameters describing the way the mice step forward were recorded, including time spent at swing, brake, propel, stride, and stance; length and frequency of strides; paw angle and its variability; and stance width. Each paw was analyzed separately. Collectively, all the parameters evaluate the muscular strength in all four limbs to detect weakness.

### Open field

tests were conducted using Open Field Systems by Kinder Scientific. The mice were placed inside an open space with high density infra-red beams. The movements of the mouse were recorded automatically based on its crossing of the infra-red beams. Total travel distance, rearing frequency, and time spent at center vs. corners of the field were analyzed over the course of 30 minutes.

### Micro-CT imaging and analysis of spine morphology

Micro-computed tomography (micro-CT) skeletal scans were acquired using an Albira system (Bruker). Animals were lightly anesthetized by 3% isoflurane and placed in a prone position on the micro-CT bed. Special attention was paid to ensure that the spine was not artificially bent, and that all limbs were in a relaxed position and not pinned underneath the torso. Stacks of cross-sectional images were reconstructed with a volume resolution of 125 μm.

A custom pipeline written in Python (https://github.com/teamnbc/MouSpine) was designed to compute three-dimensional spinal trajectories using stacks of micro-CT cross sectional images. The pipeline implements a series of annotation steps as follows. A horizontal maximum intensity projection (MIP) image is first used to place a set of points along the spinal midline and to compute a spline approximation of its trajectory (**Fig. 3A**, left). In this image, the user can also register the antero-posterior position of one reference vertebra (e.g., L6). A lateral MIP is then computed using a thin vertical stripe of voxels centered around the spline, yielding an image in which the spinal canal is clearly visible from head to tail. The user can then annotate a series of vertebral landmarks (reference points) in this image. The ventral limit of intervertebral spaces appeared as robust landmarks and were used as reference points in this study (**Fig. 3A**, middle). Using the known position of the reference vertebra defined previously, the pipeline automatically associates these points to specific vertebral identities. In the present study, each reference point was associated with the vertebra immediately anterior to it (e.g., the L6 point was located at the posterior side of L6). At this stage, only the antero-posterior and dorso-ventral coordinates of reference points are defined. The final step consists in interpolating their medio-lateral coordinates using the spline calculated above, yielding a set of Cartesian coordinates describing the three-dimensional trajectory of the spine (**Fig. 3A**, right). For one mouse, the whole annotation process is typically completed in less than 2 minutes. The person who performed these annotations was blinded to the genotype of the mice.

The rest of the analysis was performed with custom code written in R (https://github.com/teamnbc/MouSpineR, using version 4.3.1), employing a right-handed reference frame in which the *x* axis points towards the nose, the *y* axis towards the left, and the *z* axis towards the dorsal side. The first step consisted in correcting for minor body misalignments (small lateral tilt and/or slightly slantwise body axis), which are inevitable when placing an animal on the micro-CT bed. For that purpose, spinal trajectories were centered and the roll and yaw rotations minimizing the sum of squared *y* coordinates were computed and applied. A few iterations of this process suppressed apparent lateral deviations visible on horizontal projection images in tilted animals, which were in fact the signature of their natural thoracic kyphosis seen at a slight angle (**Fig. 3A**, right). Individual spinal trajectories were then aligned using a second iterative process: in each iteration, the average position of reference points was calculated; then for each trajectory, the translation within the sagittal plane minimizing the sum of squared Euclidean distances of reference points relative to their average position was computed using a grid search algorithm and applied. This process rapidly converged (with less than 10 iterations) towards an optimal alignment of spinal trajectories. Spine curvature was computed as follows. Each vertebra *i* is flanked by a pair of reference points whose coordinates can be used to compute a three-dimensional unitary vector ***v_i_*** describing the orientation of its longitudinal axis. The local curvature around each vertebra *i* was calculated as the angle between the longitudinal axes of vertebrae *i−1* and *i+1*, i.e. as the arcsine of the cross product ***v_i−1_*** x ***v_i+1_*** (**Fig. 3C**). Curvature profiles were obtained by plotting the resulting curvature angles as a function of vertebral identity, with positive and negative values representing convexities and concavities, respectively.

### Statistics

All quantitative values are presented as mean ± standard deviation (S.D.) unless specified separately. Two-tailed t-tests were used for data presented in **Figs. 2 and 4**. Sidak correction for multiple comparison was used for Fig. 4D. For Fig. 3, all statistical analyses were conducted using R version 4.2.2 (R Development Core Team, 2022, https://www.r-project.org/). Linear mixed-effects model (LMM) analyses were performed for data presented in Figure 3D. Group differences were examined using LMMs fitted to the curvature angles. In the fitted models, the factors Genotype (*Sspo*^-/-^ vs *Sspo*^+/+^), Region (thoracic vs lumbar vs sacral) and Gender (male vs female), and their interaction terms were regarded as fixed effects. Cervical and caudal regions were excluded from the analysis as they were partially and variably covered by micro-CT scans depending on the positioning of the animal along the rostro-caudal axis. The mice identifier was assigned as a random (intercept) effect to account for the paired measurements by animal across the vertebral segments. Two separate LMMs were fitted, one for values from 6-week-old mice and one for values from 12-week-old mice, using restricted maximum-likelihood estimation (REML) with the function lmer in the lme4 package (v1.1-31) [17]. For each model, significance of the main effects and interaction terms was assessed based on Type II Wald chi-square tests using the function Anova in the car package (v3.1-1), followed by post hoc comparisons of the two genotypes at each vertebra within each sex using the emmeans package (v1.4.5) with False Discovery Rate (FDR) correction of p-values. The level of statistical significance was set at *p* or adjusted *p* < 0.05 for all tests.

**Fig. 2.**
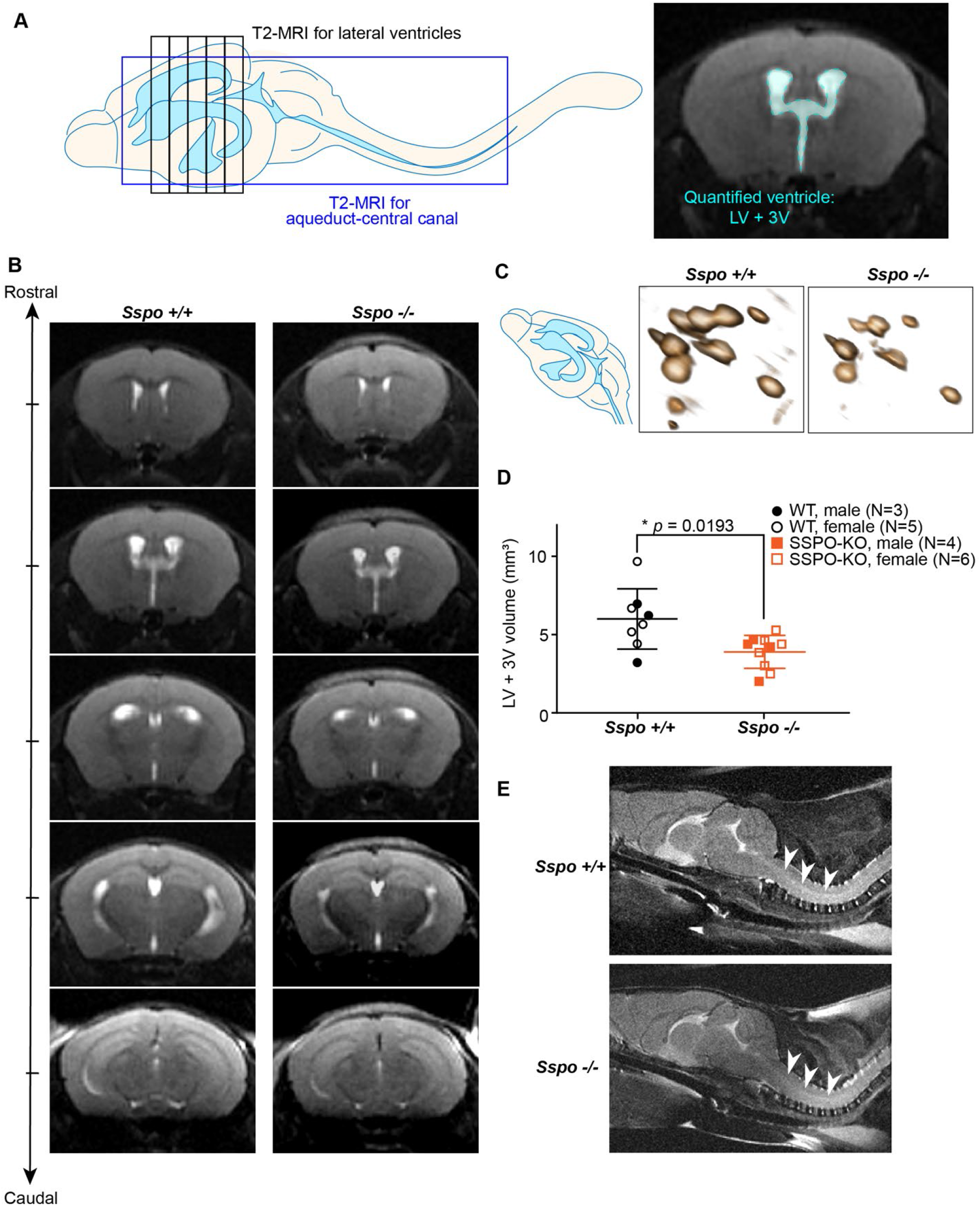
*Sspo*-deficiency is accompanied by reduced ventricle volume at 6 weeks. (**A**) Schematic depicting brain regions captured by MRI and manual ventricle segmentation. (**B**) Representative sequential MR images of *Sspo*^+/+^ and *Sspo*^-/-^ brains, showing reduced ventricle (white) sizes in *Sspo*^-/-^ mice. (**C**) 3D rendering of lateral and third ventricles from *Sspo*^+/+^ and *Sspo*^-/-^ mice. (**D**) Quantification of lateral and third ventricle volumes. Male and female mice were analyzed and plotted together but marked separately. * *p* = 0.0193, Welch’s t-test. Data presented as mean ± S.D. (**E**) Images of spinal cord showing no change in aqueduct.

**Fig. 3.**
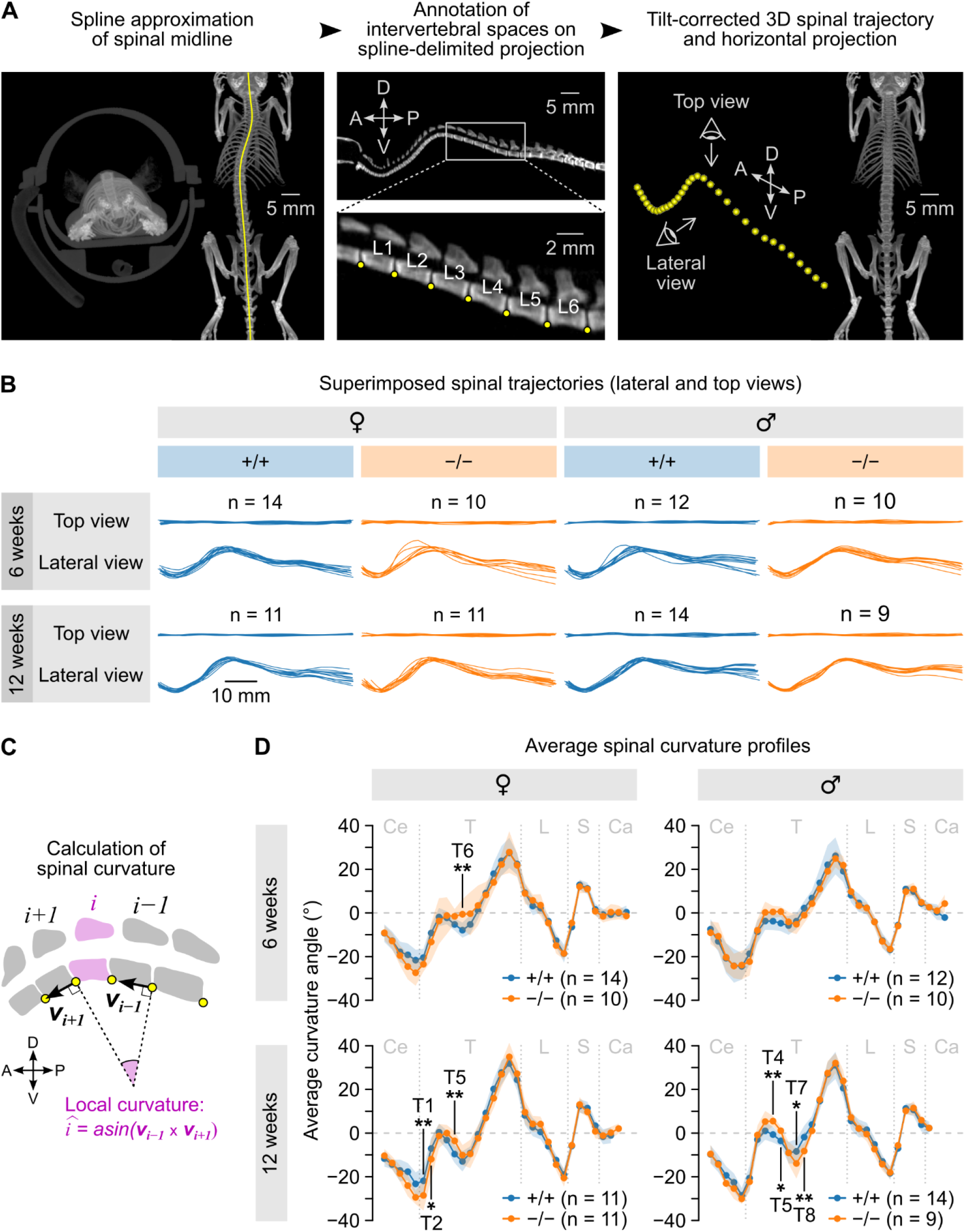
*Sspo*-deficiency only had a mild impact on spinal curvature. (**A**) Principle of the biplanar annotation approach used to measure three-dimensional spinal trajectories. A spline approximation of the spinal midline is first obtained on a horizontal projection image (right image in left panel); reference points (yellow dots) are then placed at the ventral aspect of intervertebral spaces on a lateral projection image computed from a stripe of voxels along the spline (middle); their medio-lateral position is interpolated using the spline, yielding a set of Cartesian coordinates describing the three-dimensional trajectory of the spine (right). The example mouse shown here was slightly tilted to the left, as shown by the coronal projection (left image in the left panel), resulting in an apparent deviation of the spine in the horizontal projection (right image in left panel); this apparent deviation disappeared after applying the corrective rotation computed using the three-dimensional trajectory (horizontal projection in right panel). A: anterior; D: dorsal; P: posterior; V: ventral. (**B**) Optimal alignments of individual spinal trajectories for wild type (*Sspo*^+/+^) and *Sspo*^-/-^ mice grouped by gender and age (6 and 12 weeks). In each case, top and lateral views are shown (top and bottom curves, respectively). (**C**) Sketch detailing the calculation of local curvature angles. (**D**) Average curvature angle (± S.D., shaded area) at each vertebral position computed across mice grouped by gender, age (6 and 12 weeks) and genotype. Vertebral levels at which significant differences between *Sspo*^-/-^ and *Sspo*^+/+^ mice were found using linear mixed-effects models followed by post hoc pairwise comparisons are indicated (*: *p* < 0.05; **: p < 0.001; see values in the text). Vertical gray dashed lines indicate regional limits (Ce: cervical; T: thoracic; L: lumbar; S: sacral; Ca: caudal).

**Fig. 4.**
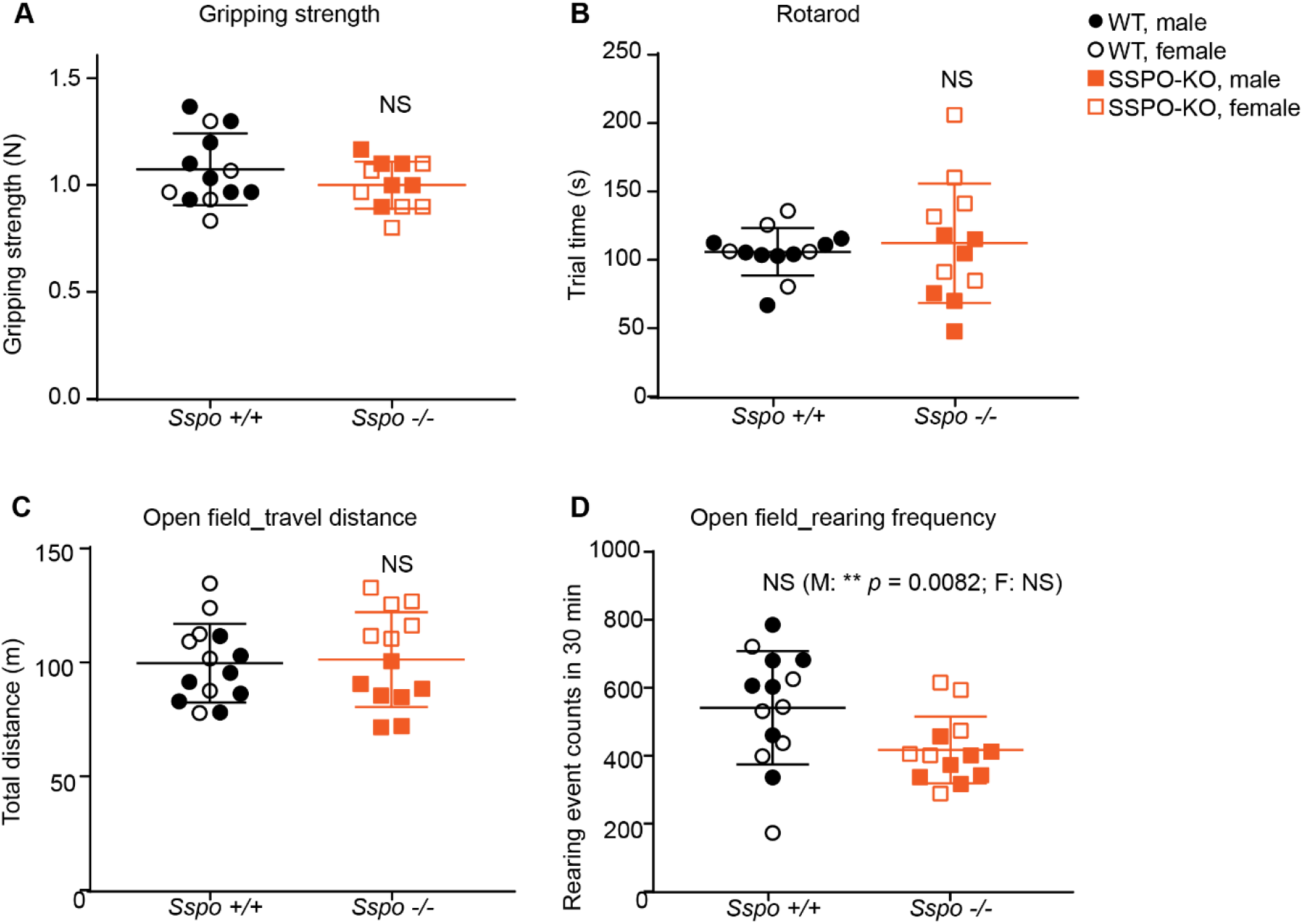
*Sspo*-deficiency did not alter typical motor behaviors at 12-14 weeks. No behavioral differences were observed in *Sspo*^-/-^ mice compared to wild type littermate controls in (**A**) forelimb gripping strength (not significant, N = 13 wild type, 8 males and 5 females, N = 12 *Sspo*^-/-^, 6 males and 6 females, *p* = 0.20), (**B**) average time spent on the rotarod (not significant, N = 13 wild type, 8 males and 5 females, N = 12 *Sspo*^-/-^, 6 males and 6 females, *p* = 0.64), (**C**) total distance traveled during each 30 min (not significant, N = 14 wild type, 7 males and 7 females, N = 13 *Sspo*^-/-^, 7 males and 6 females, *p* = 0.83). open-field test, and (**D**) number of rearing events during each 30 min (N = 14 wild type, 7 males and 7 females, N = 13 *Sspo*^-/-^, 7 males and 6 females, *p =* 0.0262 (significant *p* with Sidak correction for multiple comparison in open field test is 0.0170). If analyzing males only, ** *p* = 0.0082). open field test. All panels have male and female marked separately and analyzed together unless specified. Welch’s t-test.

## Results

### Reissner’s fiber is absent in *Sspo* knockout mice

To study the function of RF during mammalian development, we generated an *Sspo* knockout mouse with the goal of eliminating its protein product, which aggregates to form the RF. Taking a genome editing approach, four guide RNAs were designed to target exons 2, 3, 4, and 6 of *Sspo*. Of all the mutations identified in first generation progeny, one harbored a 5 bp deletion in exon 2 that shifted the open reading frame, resulting in early stop codon (**Fig. 1A**). We established the *Sspo^tm1^* mouse strain from this founder, and we refer to it as *Sspo*^-/-^.

We validated loss of *Sspo* mRNA expression in the SCO of *Sspo*^-/-^ mice using BaseScope technology. BaseScope enables the sensitive detection of very small insertions or deletions of nucleotides in genomic DNA. BaseScope probes were designed to recognize the 5 bp deletion and to distinguish between wild type (*Sspo*^+/+^, successful binding) vs. KO (*Sspo*^-/-^, failed binding) samples. Brain sections from wild type littermates showed strong *Sspo* signal in the SCO region, while no signal was detected in the SCO region or ventricles of *Sspo*^-/-^ mice (**Fig. 1B-C**). The absence of immunoreactivity against Reissner fiber material in the SCO of *Sspo*^-/-^ mice (**Fig. 1D-E**) further confirms the lack of functional *Sspo* transcripts. Importantly, while fibrous RF-positive structures were detected in wild type tissue, corresponding to loosely arranged bundles of thin filaments referred to as pre-RF material [18, 19], none was detected in *Sspo*^-/-^ mice (**Fig. 1E**, far right panels), indicating that no RF forms.

### *Sspo-*deficiency reduced brain ventricle volumes

We obtained brain scans by T2 weighted magnetic resonance imaging (MRI) in live *Sspo*^+/+^ vs. *Sspo*^-/-^ mice at 6 weeks. MR images were segmented manually by researchers blinded from the animal IDs to calculate lateral ventricle volumes (**Fig. 2A**). We analyzed a total of 8 wild type *Sspo*^+/+^ mice (3 males and 5 females) and 10 *Sspo*^-/-^ mice (4 males and 6 females). We found that *Sspo*^-/-^ mice had on average a ∼35% reduction in lateral and third ventricle size than *Sspo*^+/+^ littermate controls (**Fig. 2B-D**). These data were consistent across male and female mice and suggest that the absence of SSPO expression and RF formation during development limited lateral ventricle expansion. No structural differences in the spinal cord or central canal thickness were detected by MRI (**Fig. 2E**). The overall brain sizes were also not significantly different as measured by the average brain area obtained from cross sections of MR images (*Sspo*^+/+^: 41.71 ± 2.44 mm^2^ vs. *Sspo*^-/-^: 43.13 ± 4.24 mm^2^, *p* = 0.4146, Welch’s t-test), suggesting the smaller ventricles were not due to a smaller brain phenotype.

### Sspo-deficiency caused minor defects in the spinal geometry

To test if, as described in zebrafish [11, 20], *Sspo*^-/-^ leads to abnormal spine morphology in mice, we acquired whole-body micro-CT scans from *Sspo*^-/-^ mice and their *Sspo*^+/+^ littermates at 6 and 12 weeks of age. Because these scans did not reveal obvious spine deformation at first glance, we developed a custom morphometric analysis pipeline (see Methods) to precisely estimate spinal geometry (**Fig. 3A**). By mapping the ventral aspect of intervertebral spaces along the spinal midline (hereafter designated as “reference points”), we obtained an accurate description of three-dimensional spinal trajectories. After correcting for minor misalignments related to body placement in the micro-CT tube (**Fig. 3A**), optimal alignments of spinal trajectories were computed and mean spinal trajectories were obtained by calculating the average position of reference points. Spinal trajectories clearly varied with age and gender but did not appear to differ notably between genotypes (**Fig. 3B, Fig. S1A**). A quantitative comparison of spinal geometries was performed by computing spinal curvature at each vertebral level (see Methods and **Fig. 3C**). The influence of three factors (genotype, spinal region and gender) on curvature angles was assessed independently for 6- and 12-week-old animals using linear mixed-effects models (see Methods). As expected from average spinal trajectories (**Fig. S1**), we found a significant interaction effect of spinal region and gender at both ages (*p* = 0.030 at 6 weeks and *p* < 2.2e-16 at 12 weeks, Type II Wald chi-square test). At 6 weeks, the model taking into account all three factors only showed a non-significant trend for a three-way interaction (p = 0.063, Type II Wald chi-square test) with a post hoc comparison indicating a difference in genotypes at the level of T6 in females (*p* = 0.004, see **Supplementary File 1** – detailed statistical report, **Fig. 3D**). At 12 weeks, a significant interaction effect of all three factors was found (*p* = 0.002, Type II Wald chi-square test) and the post hoc comparison of genotypes revealed differences at the levels of T1, T2 and T5 in females (*p* = 0.003, 0.043 and 0.007, respectively, **Fig. 3D**) and at the levels of T4, T5, T7 and T8 in males (*p* = 0.004, 0.048, 0.013 and 0.005, respectively, **Fig. 3D**). These results suggest that *Sspo*^-/-^ mice tend to build up abnormal spine curvature with age in the anterior thoracic region (T1–T8). Quantitatively, this effect can be considered as very mild since differences between average curvature angles of *Sspo*^-/-^ vs *Sspo*^+/+^ mice at these levels did not exceed 7.3 degrees.

### *Sspo*-deficiency did not impact basic motor behavior

*Sspo*-deficient mice retained many motor behaviors similar to controls. Male and female mice that were 12–14 weeks old were subjected to behavioral testing to determine if they showed any changes in muscle function and mobility. The mice were tested for gripping strength, balancing ability on rotarod, stride specifics by Digi gait, and rearing and mobility in the open field. Overall, the *Sspo*^-/-^ mice lacked robust deficits in motor behaviors, except for mildly reduced rearing frequency in males (**Fig.4**). The lack of overt behavioral and mobility deficits in *Sspo*^-/-^ mice is consistent with their very mild spine deformation compared to controls.

## Discussion

In this study, we reported a new genetic mouse *Sspo*^-/-^ model created by CRISPR editing. Unlike zebrafish [11, 20], the *Sspo*^-/-^ mice exhibited very mild spine deformities confined to the thoracic level and no major motor phenotype. Instead, the *Sspo*^-/-^ mice had reduced brain ventricular volume, reflecting likely abnormality in CSF formation or dynamics.

SSPO effects on ventricular structure could be mediated by changes in physical properties of CSF such as decreased protein concentration and/or osmolarity that could lead to reduced CSF volume. CSF composition naturally changes as the brain matures, with some of the most striking changes observed in dilution of the overall CSF proteome including availability of certain health and growth-promoting factors [21] as well as changes in CSF-ion levels (e.g., K^+^) [16]. Much of the CSF protein content has been attributed to the choroid plexus. However, any cell / tissue with access to the CSF including the SCO can contribute to CSF composition. Indeed, the earliest reported presence of soluble SSPO in mouse is embryonic day 14 (E14) [22], well after neural tube closure (approx. E8) and the formation of the ventricular system. *Sspo*-deficiency could contribute to limited ventricle expansion attributed to osmolarity-mediated effects of early development [23]. Intriguingly, studies in chicken embryos suggest that when SSPO is first secreted by the SCO into the CSF in its soluble form, it has neurogenic properties and contributes to neurogenesis [24]. In our present study, we did not observe any overt changes in mouse brain size such as microcephaly.

*Sspo*^-/-^ mice had smaller ventricles at 6-weeks of age compared to wild type control and no observable abnormalities in the cerebral aqueduct, demonstrating that absence of RF alone is not sufficient to cause hydrocephalus or aqueductal stenosis. Our data contrast prior reports where SCO damage or malformation and lack of RF were reported in spontaneous hydrocephalus from aqueductal stenosis in mice and rats [25–30]. SCO damage was also reported in 2 cases of human fetal hydrocephalus [31, 32]. Because these studies observed SCO and RF changes after hydrocephalus had developed, it is possible that RF changes can arise secondary to primary injuries of stenosis, hydrocephalus, and SCO damage. One additional prior study in rats reported that immunization of a dam with RF antibodies resulted in transfer of antibodies to embryos, therefore disrupting RF formation and causing stenosis and hydrocephalus [33, 34]. Additional studies should elucidate the mechanisms underlying these different outcomes regarding SCO damage, RF formation, and ventricle expansion.

The mechanisms regulating ventricle formation and CSF volume are complex and only beginning to be understood [35, 36]. One remarkable property of SSPO is its ability to bind with numerous CSF proteins (reviewed in [2]). One can speculate that SSPO and these numerous binding partners regulate CSF osmolarity or/and secretion per se from the SCO, the ChP, and/or other paraventricular structures. Future studies will decipher between these putative interpretations.

Zebrafish *Sspo* knockouts that consistently show a curled down embryonic phenotype [11] that persists to juvenile/adult stages [20, 37, 38] and resembles defects observed in mutants with defective cilia [39–42] and/or signaling of the urotensin pathway [37, 42–44]. In contrast, *Sspo*-deficiency had a very small impact on thoracic curvature in mice. This difference may reflect species-specific mechanisms regulating spine geometry. In particular, four-limbed animals rely heavily on the proprioceptive system conveying mechanosensory feedback from the periphery for maintaining a proper spinal alignment throughout development [45, 46]. The corresponding critical contribution of dorsal root ganglia might explain why their spinal geometry would be less sensitive to perturbations affecting the interoceptive mechanosensory system constituted of the RF and the cerebrospinal fluid-contacting neurons [47–50] [58]. To characterize the mild skeletal phenotype of *Sspo^-/-^* mice, we designed a new pipeline for the quantitative analysis of spinal shape using rodent micro-CT data. Existing methods could quantify strong deformations in various mutant mouse strains [46, 51] but are not suited to detect subtle changes or thoroughly examine spinal shape. Previous solutions were usually a transposition of traditional clinical methodologies, involving the measurement of Cobb angles (the angle between the end vertebrae of an identified curved segment) using manual or semi-automated annotations of X-ray images [52]. Efforts for improving the measurement of spinal shape in rodents have remained limited to the processing of two-dimensional X-ray data [53] and therefore did not address artefactual effects due to variable animal placement (see **Fig. 3A**). Possible directions for estimating three-dimensional spinal geometry in rodents include the use of generalizable annotation tools adapted to micro-CT data such as DicomAnnotator [54] or the development of 3D reconstruction methods similar to the ones used by clinicians [55–57]. We opted for a custom annotation-based solution tailored to our specific need, i.e., estimating three-dimensional spinal trajectories without reconstructing individual vertebrae. One key element of our strategy is the calculation of a side projection image obtained from a stripe of voxels centered on the spinal midline, which greatly facilitates the visualization of vertebral landmarks such as the spinal canal and intervertebral spaces. Our method represents a tradeoff between simplicity of use (high), rapidity of development (high), complexity of the measure (low) and accuracy of the measurement (high). Our code is openly available (https://github.com/teamnbc/MouSpine) and will be adaptable to other studies with some adjustments to work with multiple data formats and non-prone positions. The companion code for performing the necessary corrections, aligning individual trajectories, computing average trajectories and calculating curvature angles is also available (https://github.com/teamnbc/MouSpineR).

Consistent with mild anatomical phenotype, function (e.g., locomotion) in the *Sspo*^-/-^ mice was minimally affected. In the open field, rotarod, and Digi gait tests, *Sspo*^-/-^ mice did not show significant differences than their wild type littermates. Their gripping strength also appeared normal. It remains possible that some mild phenotypes exist and may therefore require a finer motor analysis as observed for mice challenged in fine skilled locomotion in which CSF-cNs were ablated [48, 49]. However, these data underscore the relative adaptability of the ventricles to sustain moderate volume changes without overt disruption in brain functions. Indeed, in the clinical setting, many children with mild, non-progressive ventriculomegaly without neurodevelopmental defects will not present with long-term neurological effects.

## Conclusions

In conclusion, we created a new genetic mouse *Sspo*^-/-^ model which showed reduced brain lateral ventricle volume, implicating SSPO and RF contribution to early life CSF formation and dynamics. Unlike in zebrafish, deletion of SSPO only caused very mild spine deformation in mice at the thoracic level. Our findings underscore differences in spine morphogenesis regulation and the involvement of RF between zebrafish and limbed mammalian species like mice in which proprioceptive information from peripheral mechanoreceptors in the limbs may well play a larger role.

## List of abbreviations

CSF: cerebrospinal fluid
CSF-cN: cerebrospinal fluid contacting neurons
RF: Reissner’s fiber
SCO: Subcommissural organ
SSPO: SCO-spondin

## Declarations

### Ethics approval

All experiments involving mice in this study were approved by the BCH IACUC.

### Consent for publication

Not applicable.

### Availability of data and materials

Novel reagents are available from the corresponding author or a designated repository. The datasets used and/or analyzed during the current study are available from the corresponding author on reasonable request and made available on Dryad:

All code is available on github:

https://github.com/teamnbc/MouSpine

https://github.com/teamnbc/MouSpineR

### Authors’ contributions

H.X. generated transgenic *Sspo*^-/-^ mice, analyzed MR imaging data, performed microCT, dissected spinal cord and brains, coordinated behavioral studies; G.D. analyzed spinal geometry in *Sspo*^-/-^ and control mice; Y.C.B. performed IHC for the Reissner material on brain sections; F-X.L. implemented linear mixed models to analyze spine deformations; S.G. validated *Sspo* mRNA expression; C.W. and M.K.L. acquired funding, conceived the project, supervised research and writing of the manuscript.

## Acknowledgements

We thank members of the Lehtinen and Wyart labs for helpful discussions, Nancy Chamberlin for critical reading and editing of the manuscript, Morgan Shannon and Aja Pragana for animal husbandry support, Mantu Bhaumik and the IDDRC Gene Manipulation Core; Nathaniel Hodgson and the Animal Behavior and Physiology Core; Peter Morris, Kristina Pelkola, Simon Warfield and the BCH Small Animal Imaging Laboratory for assistance with MRI and CT; BCH IDDRC Cellular Imaging Core; Annick Prigent and the staff of the histology facility of the Paris Brain Institute (ICM).

---

**Supplementary Fig. 1.**
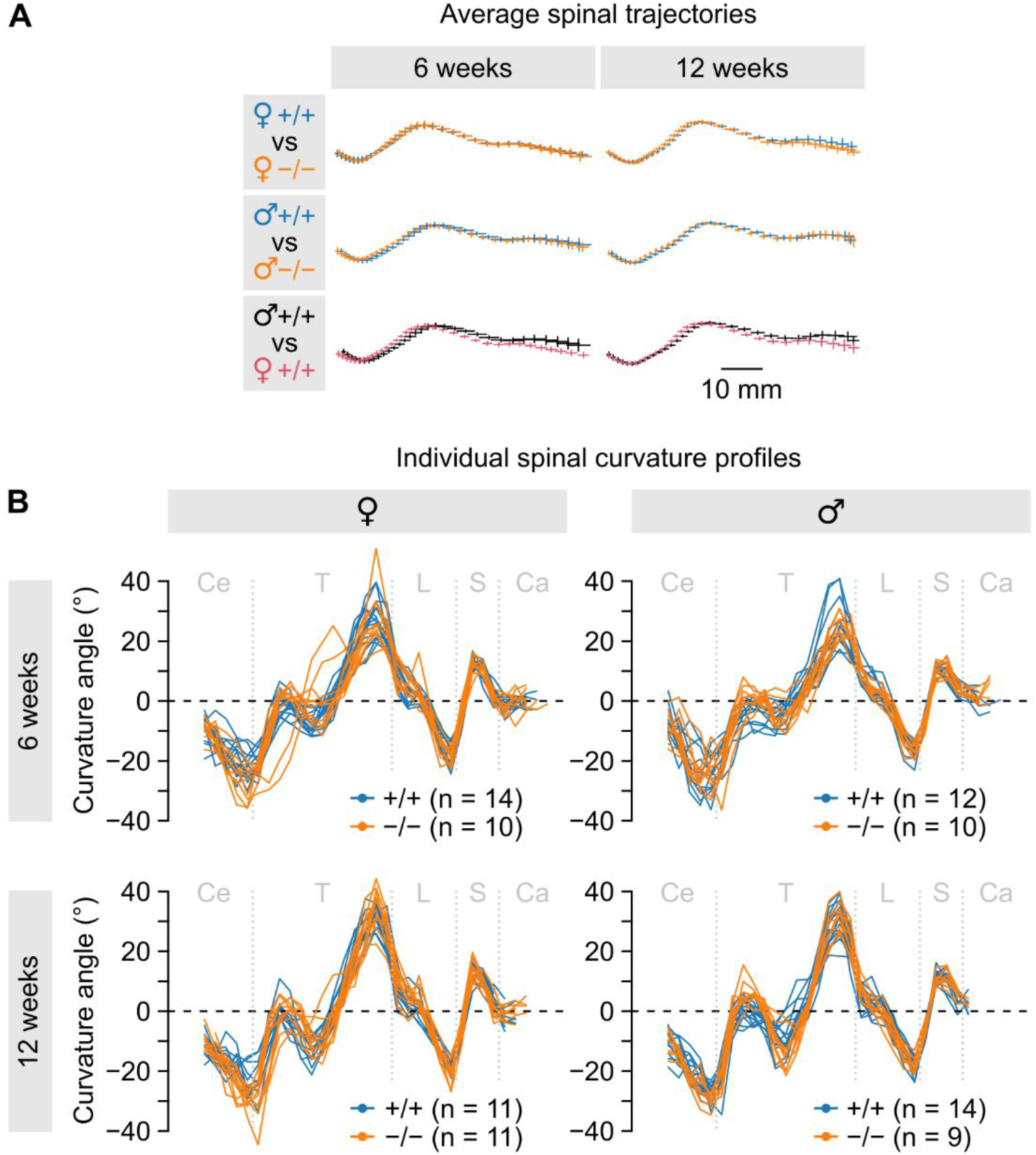
Additional information on the analysis of micro-CT data. (**A**) Optimal alignment of average spinal trajectories computed for different mouse groups. For each trajectory, the mean (*x*, *z*) position of reference points is marked using a cross whose horizontal and vertical bars represent the S.D. along *x* and *z*, respectively. (**B**) Superimposed individual curvature profiles for all mouse groups.

## Statistical analysis

## 1 Method

All statistical analyses were conducted using R version 4.2.2 (R Development Core Team, 2022) and plots were generated with the ggplot2 package (v3.4.2) (Wickham, 2016).

Differences in the angle of curvature were investigated for the two genotypes across the regions of the spine. Group differences were examined using a linear mixed-effects models (LMMs) fitted to the angle values. In the fitted models, the factors Genotype (-/-, +/+), Region (S1-S4, L1-L6, T1-T13) and Sex (M, F), and their interaction terms were regarded as fixed effects. The mice identifier was assigned as a random (intercept) effect to account for the paired measurements by animal across the spine locations. Two separate LMMs were fitted: one for the 6 weeks values and one for the 12 weeks values using restricted maximum-likelihood estimation (REML) with the function lmer in the lme4 package (Bates et al., 2015) (v1.1-31). For each model, significance of the main effects and interaction terms between Genotype, Region and Sex was assessed based on Type II Wald chi-square tests using the function Anova in the car package (v3.1-1), followed by posthoc comparisons of the two genotypes at each region of the spine within each sex using the emmeans package (v1.4.5) with False Discovery Rate (FDR) correction of P-values.

The level of statistical significance was set at *p* or adjusted *p* < 0.05 for all tests.

## 2 Data

Number of missing values. Withdraw “Ca5”, “Ca4”, “Ca3”, “Ca2”, “Ca1”, “Ce2” and “Ce1” as they contain a high number of missing values.

|  | Ca5 | Ca4 | Ca3 | Ca2 | Ca1 | S4 | Ce3 | Ce2 | Ce1 |
| --- | --- | --- | --- | --- | --- | --- | --- | --- | --- |
| F_6w | 23 | 19 | 3 | 0 | 0 | 0 | 0 | 24 | 24 |
| M_6w | 22 | 21 | 14 | 6 | 0 | 0 | 0 | 22 | 22 |
| F_12w | 22 | 22 | 18 | 9 | 1 | 0 | 1 | 22 | 22 |
| M_12w | 23 | 23 | 23 | 23 | 13 | 5 | 0 | 23 | 23 |

## 3 Plot

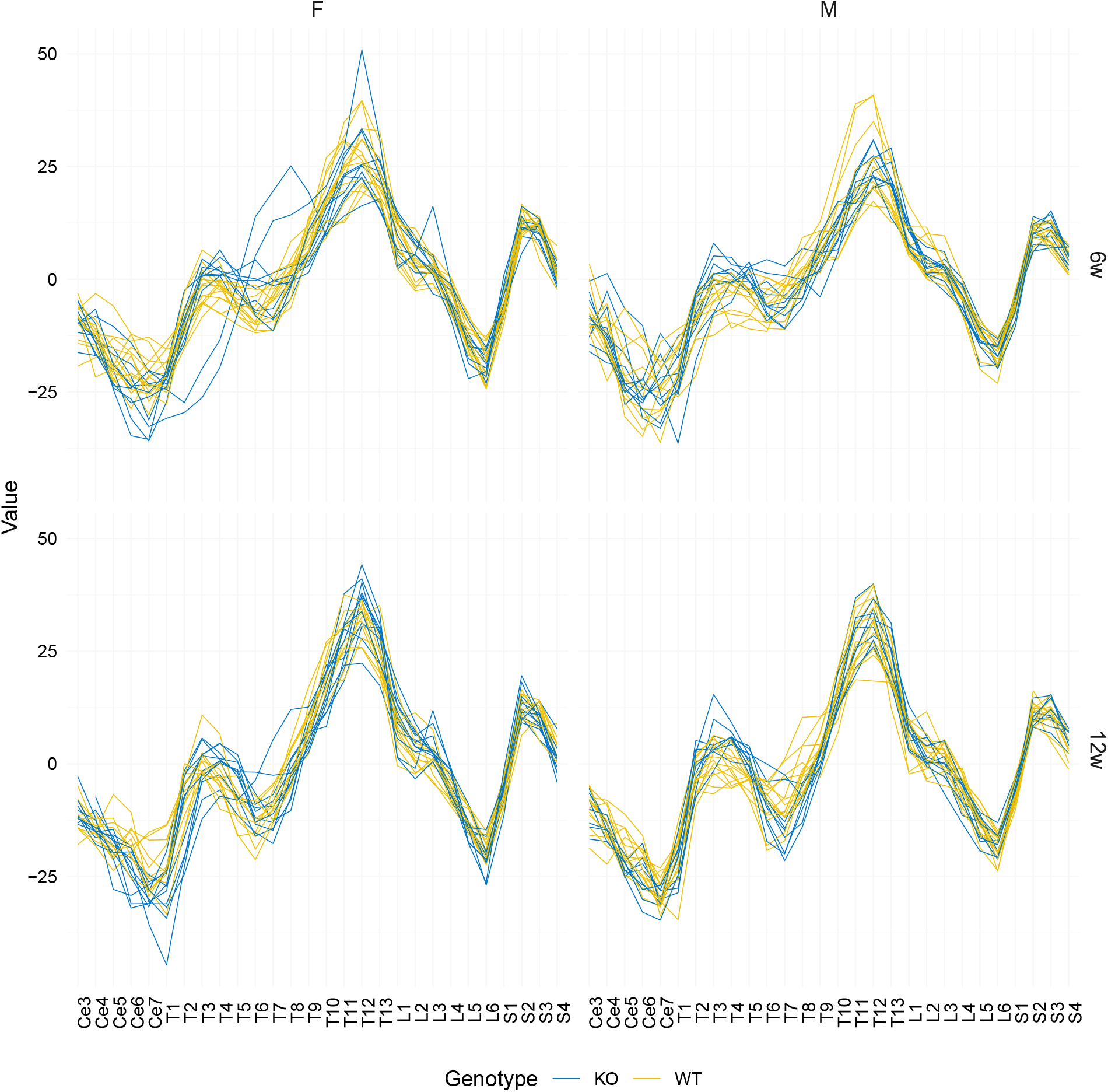

## 4 LMM 6 weeks

### 4.1 Type II Wald Chi-square tests

Type II Wald chi-square test revealed a non-significant trend towards an interaction effect of Genotype, Region and Sex on the angle of curvature (*χ*^2^(27) = 39.0, *p* = 0.063).

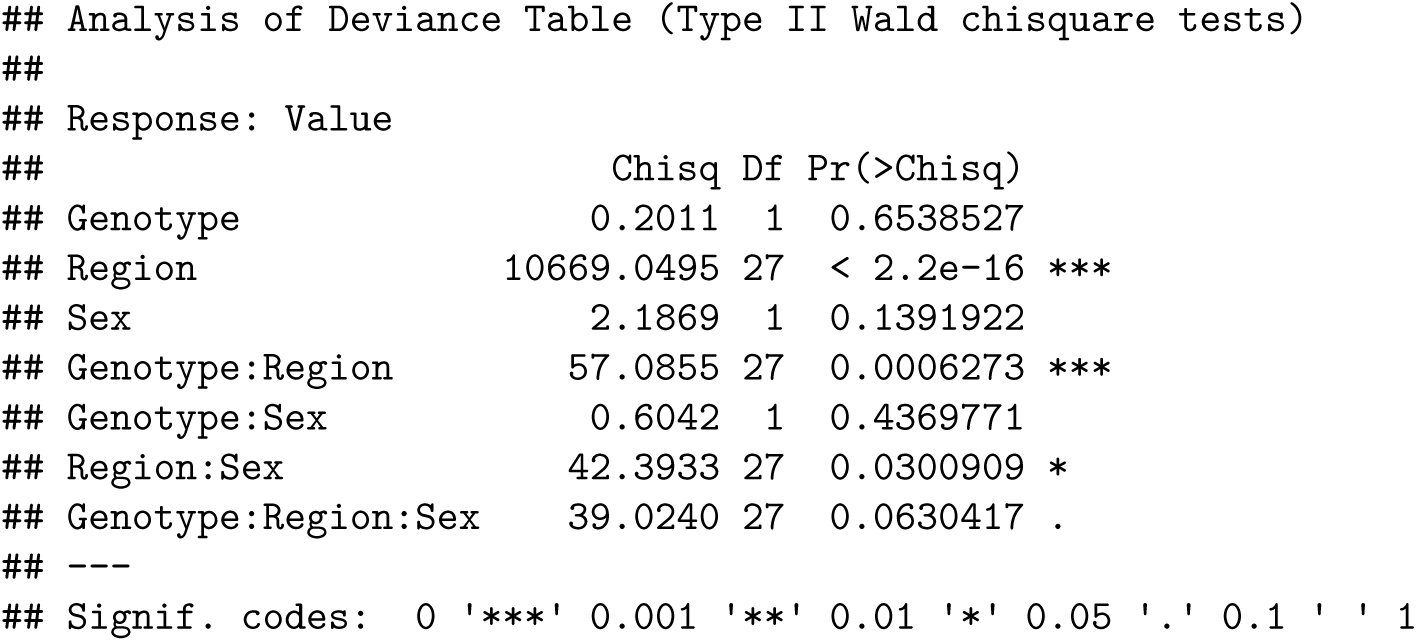

### 4.2 Posthoc emmeans comparisons

#### 4.2.1 Female - 6 weeks

Difference at T6 (*p* = 0.004).

**Table 2:** Female - 6 weeks.

| contrast | Region | estimate | SE | df | t.ratio | p.value | padj |
| --- | --- | --- | --- | --- | --- | --- | --- |
| KO - WT | T1 | -3.100 | 1.932 | 1176 | -1.604 | 0.10894 | 0.41760 |
| KO - WT | T2 | -4.099 | 1.932 | 1176 | -2.121 | 0.03410 | 0.22580 |
| KO - WT | T3 | -1.769 | 1.932 | 1176 | -0.916 | 0.36002 | 0.72471 |
| KO - WT | T4 | 1.004 | 1.932 | 1176 | 0.520 | 0.60333 | 0.85877 |
| KO - WT | T5 | 3.755 | 1.932 | 1176 | 1.943 | 0.05223 | 0.24026 |
| KO - WT | T6 | 7.288 | 1.932 | 1176 | 3.772 | 0.00017 | 0.00391 |
| KO - WT | T7 | 4.909 | 1.932 | 1176 | 2.541 | 0.01119 | 0.12868 |
| KO - WT | T8 | 2.346 | 1.932 | 1176 | 1.214 | 0.22501 | 0.62156 |
| KO - WT | T9 | -0.887 | 1.932 | 1176 | -0.459 | 0.64624 | 0.85877 |
| KO - WT | T10 | -3.988 | 1.932 | 1176 | -2.064 | 0.03927 | 0.22580 |
| KO - WT | T11 | -1.704 | 1.932 | 1176 | -0.882 | 0.37811 | 0.72471 |
| KO - WT | T12 | 0.339 | 1.932 | 1176 | 0.176 | 0.86058 | 0.91607 |
| KO - WT | T13 | -0.591 | 1.932 | 1176 | -0.306 | 0.75961 | 0.91607 |
| KO - WT | L1 | -0.318 | 1.932 | 1176 | -0.165 | 0.86925 | 0.91607 |
| KO - WT | L2 | 2.588 | 1.932 | 1176 | 1.339 | 0.18068 | 0.59366 |
| KO - WT | L3 | 1.320 | 1.932 | 1176 | 0.683 | 0.49471 | 0.85877 |
| KO - WT | L4 | -0.846 | 1.932 | 1176 | -0.438 | 0.66161 | 0.85877 |
| KO - WT | L5 | -2.256 | 1.932 | 1176 | -1.168 | 0.24322 | 0.62156 |
| KO - WT | L6 | 0.093 | 1.932 | 1176 | 0.048 | 0.96180 | 0.96180 |
| KO - WT | S1 | 1.840 | 1.932 | 1176 | 0.952 | 0.34105 | 0.72471 |
| KO - WT | S2 | -0.818 | 1.932 | 1176 | -0.423 | 0.67208 | 0.85877 |
| KO - WT | S3 | -0.301 | 1.932 | 1176 | -0.156 | 0.87624 | 0.91607 |
| KO - WT | S4 | -0.826 | 1.932 | 1176 | -0.427 | 0.66916 | 0.85877 |

#### 4.2.2 Male - 6 weeks

No difference.

**Table 3:**
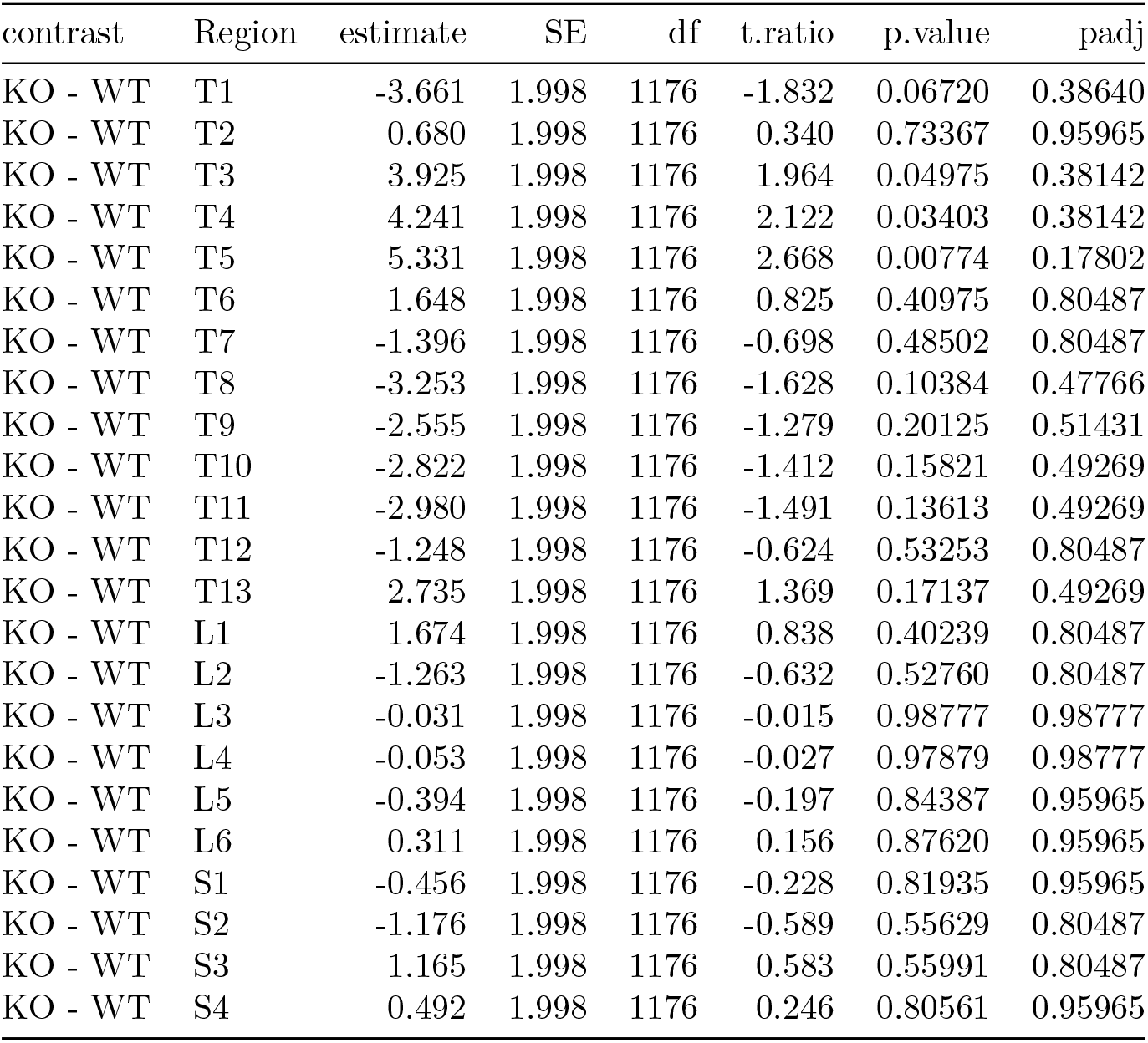
Male - 6 weeks.

### 4.3 QC

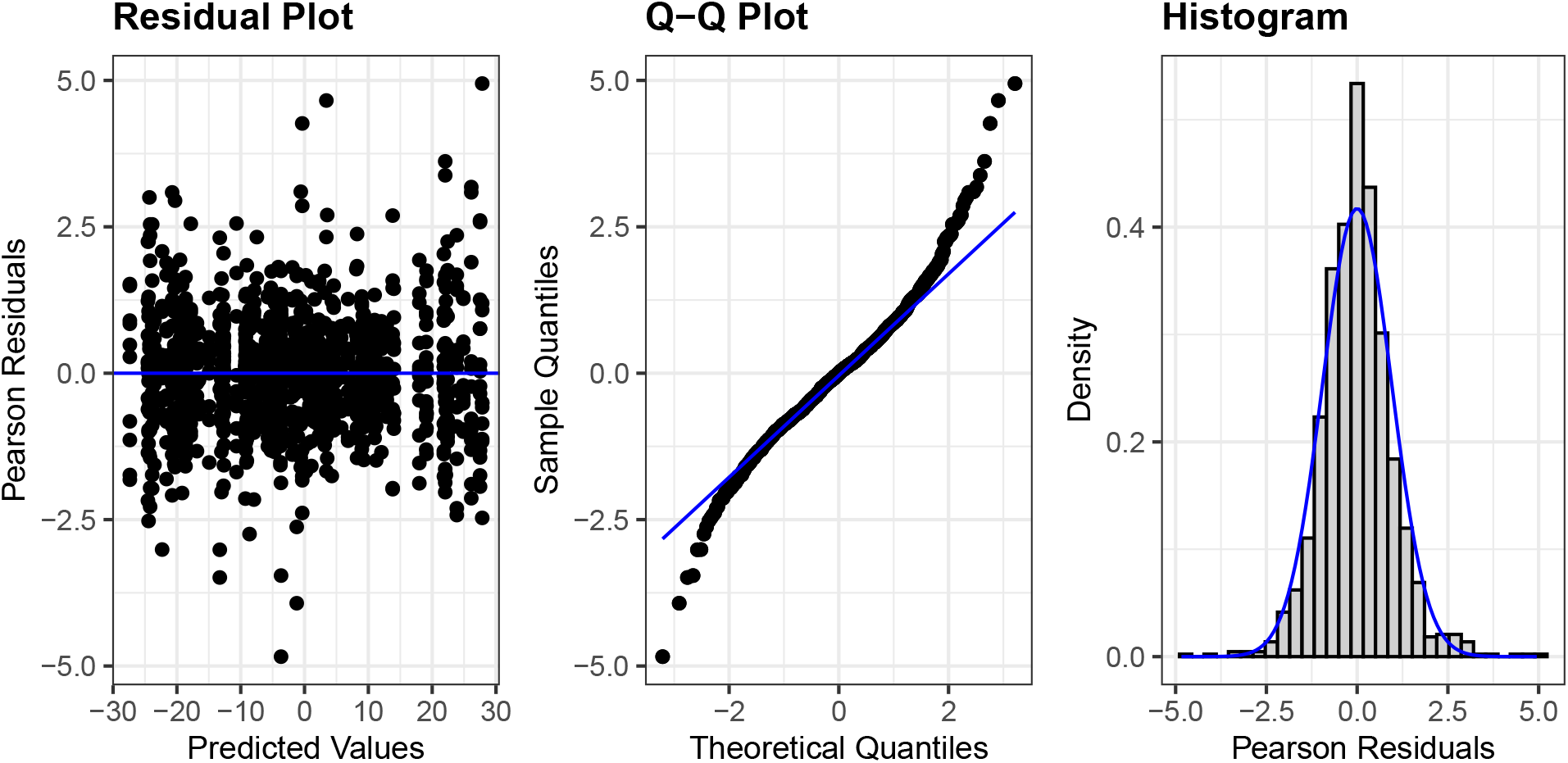

## 5 LMM 12 weeks

### 5.1 Type II Wald Chi-square tests

Type II Wald chi-square test indicated a significant interaction effect of Genotype, Region and Sex on the angle of curvature (*χ*^2^(27) = 52.7, *p* = 0.002).

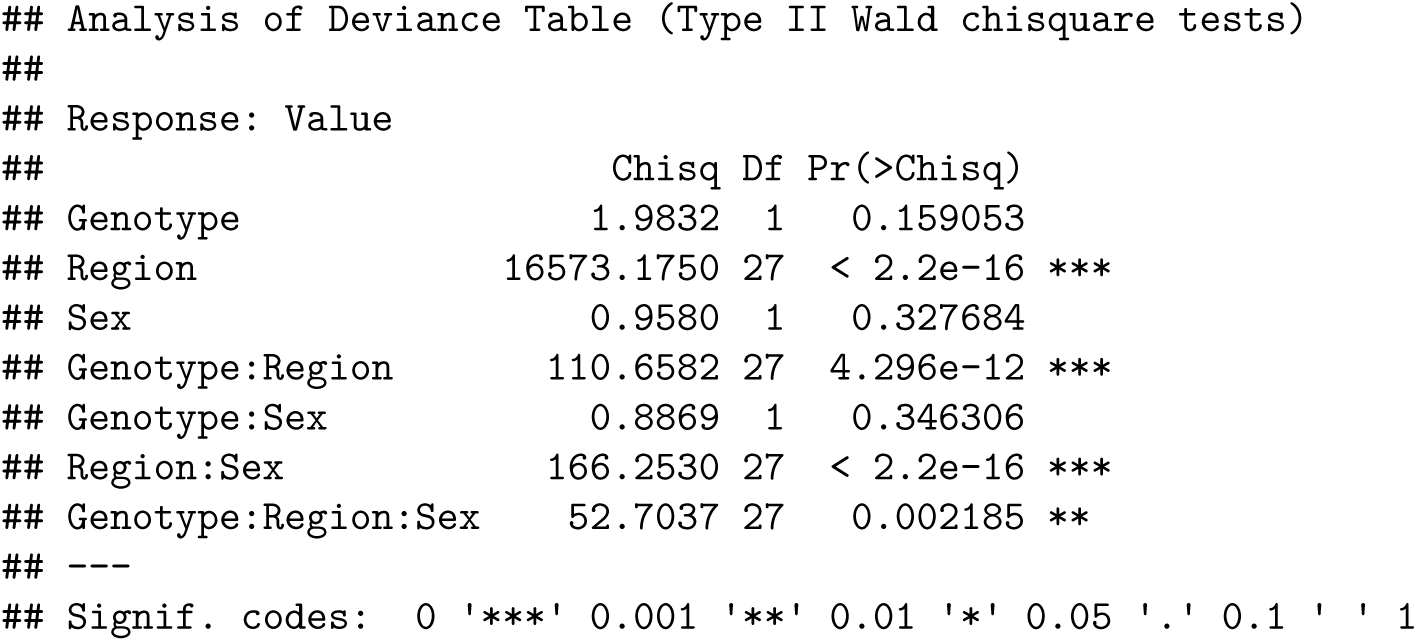

### 5.2 Posthoc emmeans comparisons

#### 5.2.1 Female - 12 weeks

Group differences at T1 (*p* = 0.003), T2 (*p* = 0.043) and T5 (*p* = 0.007).

**Table 4:** Female - 12 weeks.

| contrast | Region | estimate | SE | df | t.ratio | p.value | padj |
| --- | --- | --- | --- | --- | --- | --- | --- |
| KO - WT | T1 | -6.719 | 1.757 | 1142 | -3.825 | 0.00014 | 0.00322 |
| KO - WT | T2 | -4.873 | 1.757 | 1142 | -2.774 | 0.00563 | 0.04316 |
| KO - WT | T3 | -1.751 | 1.757 | 1142 | -0.997 | 0.31912 | 0.46578 |
| KO - WT | T4 | 3.195 | 1.757 | 1142 | 1.819 | 0.06917 | 0.26515 |
| KO - WT | T5 | 6.049 | 1.757 | 1142 | 3.443 | 0.00060 | 0.00690 |
| KO - WT | T6 | 2.872 | 1.757 | 1142 | 1.635 | 0.10238 | 0.29434 |
| KO - WT | T7 | -0.397 | 1.757 | 1142 | -0.226 | 0.82125 | 0.84939 |
| KO - WT | T8 | -1.855 | 1.757 | 1142 | -1.056 | 0.29135 | 0.46578 |
| KO - WT | T9 | -2.292 | 1.757 | 1142 | -1.305 | 0.19225 | 0.40198 |
| KO - WT | T10 | -4.224 | 1.757 | 1142 | -2.405 | 0.01634 | 0.09395 |
| KO - WT | T11 | -0.875 | 1.757 | 1142 | -0.498 | 0.61865 | 0.71145 |
| KO - WT | T12 | 3.049 | 1.757 | 1142 | 1.736 | 0.08288 | 0.27232 |
| KO - WT | T13 | 2.752 | 1.757 | 1142 | 1.567 | 0.11746 | 0.30018 |
| KO - WT | L1 | 1.489 | 1.757 | 1142 | 0.848 | 0.39684 | 0.48039 |
| KO - WT | L2 | 1.490 | 1.757 | 1142 | 0.848 | 0.39643 | 0.48039 |
| KO - WT | L3 | 3.574 | 1.757 | 1142 | 2.034 | 0.04215 | 0.19389 |
| KO - WT | L4 | 1.925 | 1.757 | 1142 | 1.096 | 0.27331 | 0.46578 |
| KO - WT | L5 | -2.070 | 1.757 | 1142 | -1.178 | 0.23889 | 0.45787 |
| KO - WT | L6 | -1.501 | 1.757 | 1142 | -0.854 | 0.39316 | 0.48039 |
| KO - WT | S1 | 0.334 | 1.757 | 1142 | 0.190 | 0.84939 | 0.84939 |
| KO - WT | S2 | 0.655 | 1.757 | 1142 | 0.373 | 0.70949 | 0.77706 |
| KO - WT | S3 | -1.733 | 1.757 | 1142 | -0.987 | 0.32402 | 0.46578 |
| KO - WT | S4 | -2.567 | 1.757 | 1142 | -1.461 | 0.14417 | 0.33159 |

#### 5.2.2 Male - 12 weeks

Differences at T4 (*p* = 0.005), T5 (*p* = 0.048), T7 (*p* = 0.014) and T8 (*p* = 0.005).

**Table 5:** Male - 12 weeks.

| contrast | Region | estimate | SE | df | t.ratio | p.value | padj |
| --- | --- | --- | --- | --- | --- | --- | --- |
| KO - WT | T1 | -1.693 | 1.760 | 1142 | -0.962 | 0.33644 | 0.64484 |
| KO - WT | T2 | 2.059 | 1.760 | 1142 | 1.170 | 0.24237 | 0.58949 |
| KO - WT | T3 | 4.244 | 1.760 | 1142 | 2.411 | 0.01605 | 0.07383 |
| KO - WT | T4 | 6.204 | 1.760 | 1142 | 3.524 | 0.00044 | 0.00506 |
| KO - WT | T5 | 4.654 | 1.760 | 1142 | 2.644 | 0.00831 | 0.04778 |
| KO - WT | T6 | -0.082 | 1.760 | 1142 | -0.047 | 0.96275 | 0.99196 |
| KO - WT | T7 | -5.513 | 1.760 | 1142 | -3.132 | 0.00178 | 0.01365 |
| KO - WT | T8 | -6.386 | 1.760 | 1142 | -3.628 | 0.00030 | 0.00506 |
| KO - WT | T9 | -2.920 | 1.760 | 1142 | -1.659 | 0.09739 | 0.32000 |
| KO - WT | T10 | 0.625 | 1.760 | 1142 | 0.355 | 0.72274 | 0.87490 |
| KO - WT | T11 | 0.483 | 1.760 | 1142 | 0.274 | 0.78389 | 0.90147 |
| KO - WT | T12 | 1.044 | 1.760 | 1142 | 0.593 | 0.55340 | 0.74872 |
| KO - WT | T13 | 3.284 | 1.760 | 1142 | 1.866 | 0.06236 | 0.23905 |
| KO - WT | L1 | 1.554 | 1.760 | 1142 | 0.883 | 0.37753 | 0.66794 |
| KO - WT | L2 | 0.018 | 1.760 | 1142 | 0.010 | 0.99196 | 0.99196 |
| KO - WT | L3 | -1.999 | 1.760 | 1142 | -1.136 | 0.25630 | 0.58949 |
| KO - WT | L4 | -2.174 | 1.760 | 1142 | -1.235 | 0.21701 | 0.58949 |
| KO - WT | L5 | -1.275 | 1.760 | 1142 | -0.724 | 0.46917 | 0.71939 |
| KO - WT | L6 | 0.326 | 1.760 | 1142 | 0.185 | 0.85308 | 0.93433 |
| KO - WT | S1 | 1.383 | 1.760 | 1142 | 0.786 | 0.43216 | 0.70998 |
| KO - WT | S2 | -1.071 | 1.760 | 1142 | -0.608 | 0.54308 | 0.74872 |
| KO - WT | S3 | 0.841 | 1.760 | 1142 | 0.478 | 0.63301 | 0.80885 |
| KO - WT | S4 | 2.090 | 1.943 | 1142 | 1.076 | 0.28214 | 0.58993 |

### 5.3 QC

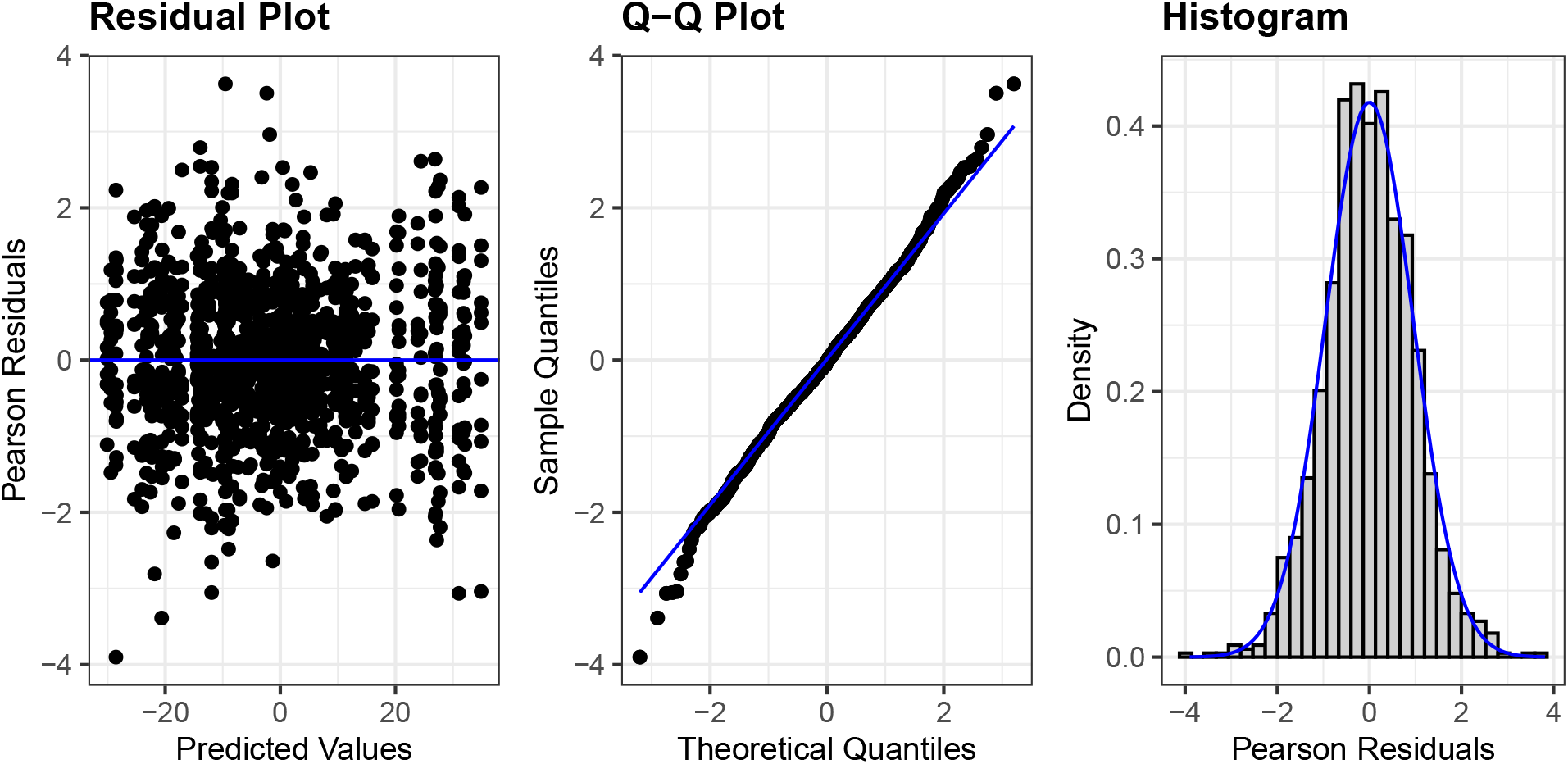

## 6 R session information

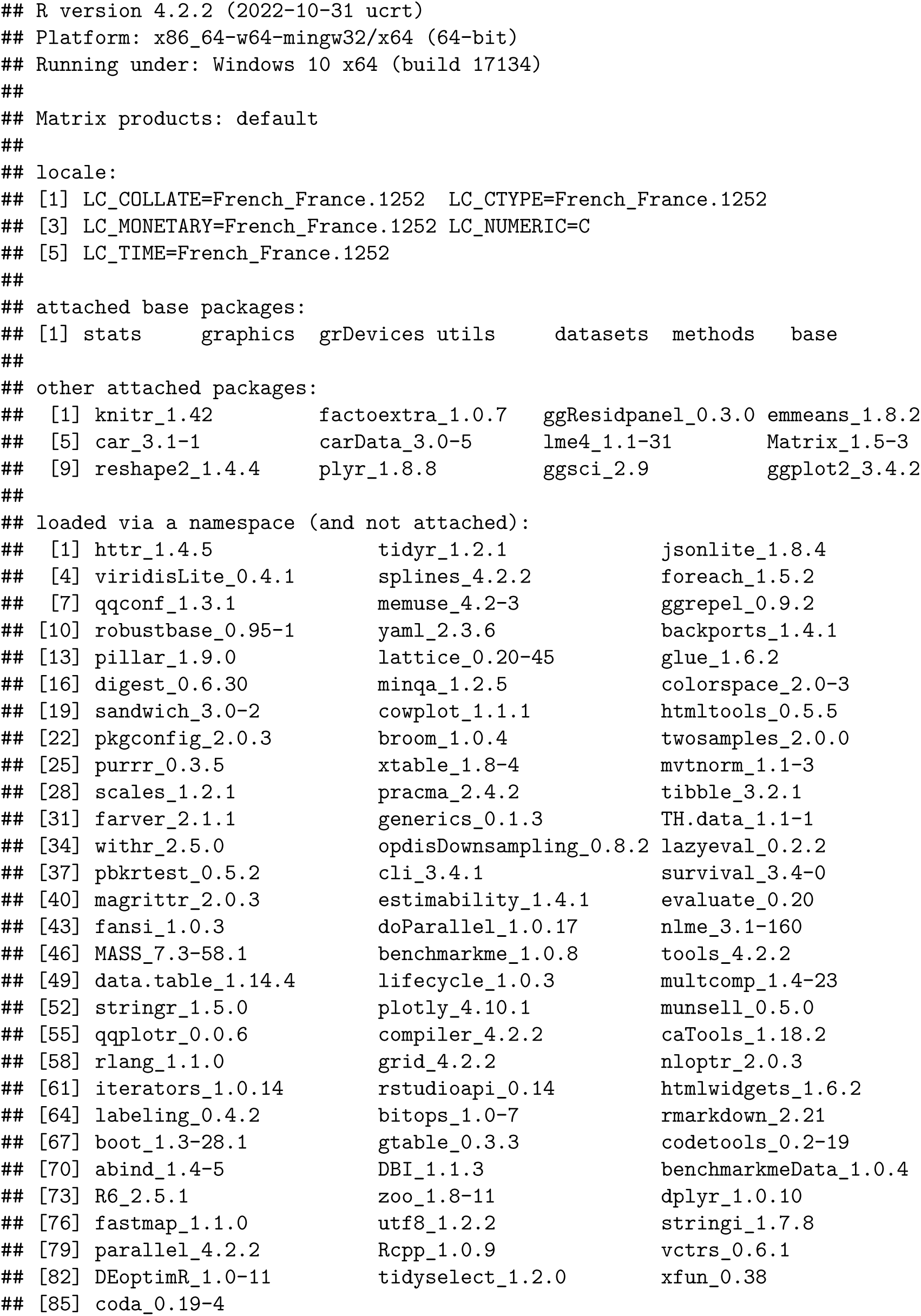

## REFERENCES

1. Montecinos, H.A., H. Richter, T. Caprile, and E.M. Rodriguez, Synthesis of transthyretin by the ependymal cells of the subcommissural organ. Cell Tissue Res, 2005. 320(3): p. 487–99

2. Sepulveda, V., F. Maurelia, M. Gonzalez, J. Aguayo, and T. Caprile, SCO-spondin, a giant matricellular protein that regulates cerebrospinal fluid activity. Fluids Barriers CNS, 2021. 18(1): p. 45.PMC8487547

3. Sterba, G., C. Kiessig, W. Naumann, H. Petter, and I. Kleim, The secretion of the subcommissural organ. A comparative immunocytochemical investigation. Cell Tissue Res, 1982. 226(2): p. 427–39

4. Gobron, S., I. Creveaux, R. Meiniel, R. Didier, A. Herbet, M. Bamdad, F. El Bitar, B. Dastugue, and A. Meiniel, Subcommissural organ/Reissner’s fiber complex: characterization of SCO-spondin, a glycoprotein with potent activity on neurite outgrowth. Glia, 2000. 32(2): p. 177–91

5. Meiniel, O. and A. Meiniel, The complex multidomain organization of SCO-spondin protein is highly conserved in mammals. Brain Res Rev, 2007. 53(2): p. 321–7

6. Gobron, S., H. Monnerie, R. Meiniel, I. Creveaux, W. Lehmann, D. Lamalle, B. Dastugue, and A. Meiniel, SCO-spondin: a new member of the thrombospondin family secreted by the subcommissural organ is a candidate in the modulation of neuronal aggregation. J Cell Sci, 1996. 109 (Pt 5): p. 1053–61

7. Guerra, M.M., C. Gonzalez, T. Caprile, M. Jara, K. Vio, R.I. Munoz, S. Rodriguez, and E.M. Rodriguez, Understanding How the Subcommissural Organ and Other Periventricular Secretory Structures Contribute via the Cerebrospinal Fluid to Neurogenesis. Front Cell Neurosci, 2015. 9: p. 480.PMC4689152

8. Sakka, L., N. Deletage, F. Lalloue, A. Duval, J. Chazal, J.J. Lemaire, A. Meiniel, H. Monnerie, and S. Gobron, SCO-spondin derived peptide NX210 induces neuroprotection in vitro and promotes fiber regrowth and functional recovery after spinal cord injury. PLoS One, 2014. 9(3): p. e93179.PMC3965545 development. S. Gobron is the Chief executive officer of Neuronax, and Nathalie Deletage and Amelie Duval are employed by Neuronax. There are no further patents, products in development or marketed products to declare. This does not alter the authors’ adherence to all the PLOS ONE policies on sharing data and materials, as detailed online in the guide for authors.

9. Vera, A., A. Recabal, N. Saldivia, K. Stanic, M. Torrejon, H. Montecinos, and T. Caprile, Interaction between SCO-spondin and low density lipoproteins from embryonic cerebrospinal fluid modulates their roles in early neurogenesis. Front Neuroanat, 2015. 9: p. 72.PMC4446542

10. Aboitiz, F. and J.F. Montiel, The Enigmatic Reissner’s Fiber and the Origin of Chordates. Front Neuroanat, 2021. 15: p. 703835.PMC8261243

11. Cantaut-Belarif, Y., J.R. Sternberg, O. Thouvenin, C. Wyart, and P.L. Bardet, The Reissner Fiber in the Cerebrospinal Fluid Controls Morphogenesis of the Body Axis. Curr Biol, 2018. 28(15): p. 2479–2486 e4.PMC6089837

12. Bagnat, M. and R.S. Gray, Development of a straight vertebrate body axis. Development, 2020. 147(21).PMC7561478

13. Bearce, E.A. and D.T. Grimes, On being the right shape: Roles for motile cilia and cerebrospinal fluid flow in body and spine morphology. Semin Cell Dev Biol, 2021. 110: p. 104–112

14. Didier, R., B. Dastugue, and A. Meiniel, The secretory material of the subcommissural organ of the chick embryo. Characterization of a specific polypeptide by two-dimensional electrophoresis. Int J Dev Biol, 1995. 39(3): p. 493–9

15. Schindelin, J., I. Arganda-Carreras, E. Frise, V. Kaynig, M. Longair, T. Pietzsch, S. Preibisch, C. Rueden, S. Saalfeld, B. Schmid, J.Y. Tinevez, D.J. White, V. Hartenstein, K. Eliceiri, P. Tomancak, and A. Cardona, Fiji: an open-source platform for biological-image analysis. Nat Methods, 2012. 9(7): p. 676–82.3855844

16. Xu, H., R.M. Fame, C. Sadegh, J. Sutin, C. Naranjo, S. Della, J. Cui, F.B. Shipley, A. Vernon, F. Gao, Y. Zhang, M.J. Holtzman, M. Heiman, B.C. Warf, P.Y. Lin, and M.K. Lehtinen, Choroid plexus NKCC1 mediates cerebrospinal fluid clearance during mouse early postnatal development. Nat Commun, 2021. 12(1): p. 447.PMC7815709

17. Bates, D., M. Mächler, B. Bolker, and S. Walker, Fitting Linear Mixed-Effects Models Using lme4. Journal of Statistical Software, 2015. 67(1): p. 1–48

18. Losecke, W., W. Naumann, and G. Sterba, Preparation and discharge of secretion in the subcommissural organ of the rat. An electron-microscopic immunocytochemical study. Cell Tissue Res, 1984. 235(1): p. 201–6

19. Rodriguez, E.M., H. Herrera, B. Peruzzo, S. Rodriguez, S. Hein, and A. Oksche, Light- and electron-microscopic immunocytochemistry and lectin histochemistry of the subcommissural organ: evidence for processing of the secretory material. Cell Tissue Res, 1986. 243(3): p. 545–59

20. Rose, C.D., D. Pompili, K. Henke, J.L.M. Van Gennip, A. Meyer-Miner, R. Rana, S. Gobron, M.P. Harris, M. Nitz, and B. Ciruna, SCO-Spondin Defects and Neuroinflammation Are Conserved Mechanisms Driving Spinal Deformity across Genetic Models of Idiopathic Scoliosis. Curr Biol, 2020. 30(12): p. 2363–2373 e6

21. Lun, M.P., M.B. Johnson, K.G. Broadbelt, M. Watanabe, Y.J. Kang, K.F. Chau, M.W. Springel, A. Malesz, A.M. Sousa, M. Pletikos, T. Adelita, M.L. Calicchio, Y. Zhang, M.J. Holtzman, H.G. Lidov, N. Sestan, H. Steen, E.S. Monuki, and M.K. Lehtinen, Spatially heterogeneous choroid plexus transcriptomes encode positional identity and contribute to regional CSF production. J Neurosci, 2015. 35(12): p. 4903–16.4389594

22. Rakic, P. and R.L. Sidman, Subcommissural organ and adjacent ependyma: autoradiographic study of their origin in the mouse brain. Am J Anat, 1968. 122(2): p. 317–35

23. Gato, A. and M.E. Desmond, Why the embryo still matters: CSF and the neuroepithelium as interdependent regulators of embryonic brain growth, morphogenesis and histiogenesis. Dev Biol, 2009. 327(2): p. 263–72

24. Vera, A., K. Stanic, H. Montecinos, M. Torrejon, S. Marcellini, and T. Caprile, SCO-spondin from embryonic cerebrospinal fluid is required for neurogenesis during early brain development. Front Cell Neurosci, 2013. 7: p. 80.PMC3669746

25. Pérez-Fígares, J.M., M. Jiménez Aj Fau - Pérez-Martín, P. Pérez-Martín M Fau - Fernández-Llebrez, M. Fernández-Llebrez P Fau - Cifuentes, P. Cifuentes M Fau - Riera, S. Riera P Fau - Rodríguez, E.M. Rodríguez S Fau - Rodríguez, and E.M. Rodríguez, Spontaneous congenital hydrocephalus in the mutant mouse hyh. Changes in the ventricular system and the subcommissural organ. (0022-3069 (Print))

26. Takeuchi, I.K., R. Kimura, M. Matsuda, and R. Shoji, Absence of subcommissural organ in the cerebral aqueduct of congenital hydrocephalus spontaneously occurring in MT/HokIdr mice. Acta Neuropathol, 1987. 73(4): p. 320–2

27. Takeuchi, I.K., R. Kimura, and R. Shoji, Dysplasia of subcommissural organ in congenital hydrocephalus spontaneously occurring in CWS/Idr rats. Experientia, 1988. 44(4): p. 338–40

28. Louvi, A. and M. Wassef, Ectopic engrailed 1 expression in the dorsal midline causes cell death, abnormal differentiation of circumventricular organs and errors in axonal pathfinding. Development, 2000. 127(18): p. 4061–71

29. Fernandez-Llebrez, P., J.M. Grondona, J. Perez, M.F. Lopez-Aranda, G. Estivill-Torrus, P.F. Llebrez-Zayas, E. Soriano, C. Ramos, Y. Lallemand, A. Bach, and B. Robert, Msx1-deficient mice fail to form prosomere 1 derivatives, subcommissural organ, and posterior commissure and develop hydrocephalus. J Neuropathol Exp Neurol, 2004. 63(6): p. 574–86

30. Ramos, C., P. Fernandez-Llebrez, A. Bach, B. Robert, and E. Soriano, Msx1 disruption leads to diencephalon defects and hydrocephalus. Dev Dyn, 2004. 230(3): p. 446–60

31. Castaneyra-Perdomo, A., G. Meyer, E. Carmona-Calero, J. Banuelos-Pineda, R. Mendez-Medina, C. Ormazabal-Ramos, and R. Ferres-Torres, Alterations of the subcommissural organ in the hydrocephalic human fetal brain. Brain Res Dev Brain Res, 1994. 79(2): p. 316–20

32. Ortega, E., R.I. Munoz, N. Luza, F. Guerra, M. Guerra, K. Vio, R. Henzi, J. Jaque, S. Rodriguez, J.P. McAllister, and E. Rodriguez, The value of early and comprehensive diagnoses in a human fetus with hydrocephalus and progressive obliteration of the aqueduct of Sylvius: Case Report. BMC Neurol, 2016. 16: p. 45.PMC4828774

33. Rodriguez, S., K. Vio, C. Wagner, M. Barria, E.H. Navarrete, V.D. Ramirez, J.M. Perez-Figares, and E.M. Rodriguez, Changes in the cerebrospinal-fluid monoamines in rats with an immunoneutralization of the subcommissural organ-Reissner’s fiber complex by maternal delivery of antibodies. Exp Brain Res, 1999. 128(3): p. 278–90

34. Vio, K., S. Rodriguez, E.H. Navarrete, J.M. Perez-Figares, A.J. Jimenez, and E.M. Rodriguez, Hydrocephalus induced by immunological blockage of the subcommissural organ-Reissner’s fiber (RF) complex by maternal transfer of anti-RF antibodies. Exp Brain Res, 2000. 135(1): p. 41–52

35. Fame, R.M., C. Cortes-Campos, and H.L. Sive, Brain Ventricular System and Cerebrospinal Fluid Development and Function: Light at the End of the Tube: A Primer with Latest Insights. Bioessays, 2020. 42(3): p. e1900186

36. Lowery, L.A. and H. Sive, Totally tubular: the mystery behind function and origin of the brain ventricular system. Bioessays, 2009. 31(4): p. 446–58.PMC3003255

37. Lu, H., A. Shagirova, J.L. Goggi, H.L. Yeo, and S. Roy, Reissner fibre-induced urotensin signalling from cerebrospinal fluid-contacting neurons prevents scoliosis of the vertebrate spine. Biol Open, 2020. 9(5).PMC7240301

38. Troutwine, B.R., P. Gontarz, M.J. Konjikusic, R. Minowa, A. Monstad-Rios, D.S. Sepich, R.Y. Kwon, L. Solnica-Krezel, and R.S. Gray, The Reissner Fiber Is Highly Dynamic In Vivo and Controls Morphogenesis of the Spine. Curr Biol, 2020. 30(12): p. 2353–2362 e3.PMC7891109

39. Christine, V., A. Isabelle, P. Guillaume, C.-B. Yasmine, E. Alexis, D. Morgane, S. Diego López, R. Hélène Le, J. Arnim, K. Hanane, V. Joëlle, P. Caroline, and S.-M. Sylvie, Loss of the Reissner Fiber and increased URP neuropeptide signaling underlie scoliosis in a zebrafish ciliopathy mutant. bioRxiv, 2019: p. 2019.12.19.882258

40. Grimes, D.T., C.W. Boswell, N.F. Morante, R.M. Henkelman, R.D. Burdine, and B. Ciruna, Zebrafish models of idiopathic scoliosis link cerebrospinal fluid flow defects to spine curvature. Science, 2016. 352(6291): p. 1341–4.PMC5574193

41. Xie, H., Y. Kang, J. Liu, M. Huang, Z. Dai, J. Shi, S. Wang, L. Li, Y. Li, P. Zheng, Y. Sun, Q. Han, J. Zhang, Z. Zhu, L. Xu, P.C. Yelick, M. Cao, and C. Zhao, Ependymal polarity defects coupled with disorganized ciliary beating drive abnormal cerebrospinal fluid flow and spine curvature in zebrafish. PLoS Biol, 2023. 21(3): p. e3002008.PMC10013924

42. Zhang, X., S. Jia, Z. Chen, Y.L. Chong, H. Xie, D. Feng, X. Wu, D.Z. Song, S. Roy, and C. Zhao, Cilia-driven cerebrospinal fluid flow directs expression of urotensin neuropeptides to straighten the vertebrate body axis. Nat Genet, 2018. 50(12): p. 1666–1673

43. Bearce, E.A., Z.H. Irons, J.R. O’Hara-Smith, C.J. Kuhns, S.I. Fisher, W.E. Crow, and D.T. Grimes, Urotensin II-related peptides, Urp1 and Urp2, control zebrafish spine morphology. Elife, 2022. 11.PMC9836392

44. Gaillard, A.L., T. Mohamad, F.B. Quan, A. de Cian, C. Mosimann, H. Tostivint, and G. Pezeron, Urp1 and Urp2 act redundantly to maintain spine shape in zebrafish larvae. Dev Biol, 2023. 496: p. 36–51

45. Blecher, R., L. Heinemann-Yerushalmi, E. Assaraf, N. Konstantin, J.R. Chapman, T.C. Cope, G.S. Bewick, R.W. Banks, and E. Zelzer, New functions for the proprioceptive system in skeletal biology. Philos Trans R Soc Lond B Biol Sci, 2018. 373(1759).PMC6158198

46. Blecher, R., S. Krief, T. Galili, I.E. Biton, T. Stern, E. Assaraf, D. Levanon, E. Appel, Y. Anekstein, G. Agar, Y. Groner, and E. Zelzer, The Proprioceptive System Masterminds Spinal Alignment: Insight into the Mechanism of Scoliosis. Dev Cell, 2017. 42(4): p. 388–399 e3

47. Orts-Del’Immagine, A., Y. Cantaut-Belarif, O. Thouvenin, J. Roussel, A. Baskaran, D. Langui, F. Koeth, P. Bivas, F.X. Lejeune, P.L. Bardet, and C. Wyart, Sensory Neurons Contacting the Cerebrospinal Fluid Require the Reissner Fiber to Detect Spinal Curvature In Vivo. Curr Biol, 2020. 30(5): p. 827–839 e4

48. Gerstmann, K., N. Jurcic, E. Blasco, S. Kunz, F. de Almeida Sassi, N. Wanaverbecq, and N. Zampieri, The role of intraspinal sensory neurons in the control of quadrupedal locomotion. Curr Biol, 2022. 32(11): p. 2442–2453 e4

49. Nakamura, Y., M. Kurabe, M. Matsumoto, T. Sato, S. Miyashita, K. Hoshina, Y. Kamiya, K. Tainaka, H. Matsuzawa, N. Ohno, and M. Ueno, Cerebrospinal fluid-contacting neuron tracing reveals structural and functional connectivity for locomotion in the mouse spinal cord. Elife, 2023. 12.PMC9943067

50. Wyart, C., Unraveling the roles of cerebrospinal fluid-contacting neurons. Elife, 2023. 12.PMC10038656

51. Gao, C., B.P. Chen, M.B. Sullivan, J. Hui, J.A. Ouellet, J.E. Henderson, and N. Saran, Micro CT Analysis of Spine Architecture in a Mouse Model of Scoliosis. Front Endocrinol (Lausanne), 2015. 6: p. 38.PMC4365746

52. Zhang, J., H. Li, L. Lv, and Y. Zhang, Computer-Aided Cobb Measurement Based on Automatic Detection of Vertebral Slopes Using Deep Neural Network. Int J Biomed Imaging, 2017. 2017: p. 9083916.PMC5651147

53. Okashi, O.A., H. Du, and H. Al-Assam, Automatic spine curvature estimation from X-ray images of a mouse model. Comput Methods Programs Biomed, 2017. 140: p. 175–184

54. Dong, Q., G. Luo, D. Haynor, M. O’Reilly, K. Linnau, Z. Yaniv, J.G. Jarvik, and N. Cross, DicomAnnotator: a Configurable Open-Source Software Program for Efficient DICOM Image Annotation. J Digit Imaging, 2020. 33(6): p. 1514–1526.PMC7728983

55. Galbusera, F., G. Casaroli, and T. Bassani, Artificial intelligence and machine learning in spine research. JOR Spine, 2019. 2(1): p. e1044.PMC6686793

56. Humbert, L., J.A. De Guise, B. Aubert, B. Godbout, and W. Skalli, 3D reconstruction of the spine from biplanar X-rays using parametric models based on transversal and longitudinal inferences. Med Eng Phys, 2009. 31(6): p. 681–7

57. Suri, A., B.C. Jones, G. Ng, N. Anabaraonye, P. Beyrer, A. Domi, G. Choi, S. Tang, A. Terry, T. Leichner, I. Fathali, N. Bastin, H. Chesnais, and C.S. Rajapakse, A deep learning system for automated, multi-modality 2D segmentation of vertebral bodies and intervertebral discs. Bone, 2021. 149: p. 115972.PMC8217255

58. Wyart, C, Carbo-Tano, M, Cantaut-Belarif, Y, Orts-Del-Immagine, A, Böhm, UL. Cerebrospinal fluid-contacting neurons: multimodal roles in the CNS. In press. https://doi.org/10.1038/s41583-023-00723-8

## References

R Core Team (2022). R: A language and environment for statistical computing. R Foundation for Statistical Computing, Vienna, Austria. URL https://www.R-project.org/.

Wickham H. 2016. Ggplot2: Elegant Graphics for Data Analysis. Springer.

Bates D, Maechler M, Bolker B, Walker S. Fitting Linear Mixed Effects Models Using lme4. J Stat Softw. 2015;67:1–48.

